# Inhibitory Cerebellar Stimulation Increases Cortical Activation: Evidence for Cerebellar Scaffolding of Cortical Processing

**DOI:** 10.1101/2021.11.02.466978

**Authors:** Ted Maldonado, T. Bryan Jackson, Jessica A. Bernard

## Abstract

While the cerebellum makes known contributions to non-motor task performance, the role of the structure remains unknown. One possibility is that the cerebellum allows for the offloading of cortical processing, providing support during task performance, using internal models. The current work used transcranial direct current stimulation to modulate cerebellar function and investigate the impact on cortical activation patterns. Participants received stimulation over the right cerebellum before a functional magnetic resonance imaging scan where participants completed a sequence learning and a working memory task. We predicted that cathodal stimulation would improve, and anodal stimulation would hinder task performance and cortical activation. We found that anodal cerebellar stimulation resulted in increased bilateral cortical activation, particularly in parietal and frontal regions known to be involved in cognitive processing. This suggests that if the cerebellum is not functioning optimally, there is a greater need for cortical resources.

## Introduction

Interest in the role the cerebellum plays in non-motor cognitive processing has increased over the last 30 years ^1^. In addition to cerebellar lesion work (Ilg et al., 2013; Schmahmann & Sherman, 1998; Timmann et al., 2008), imaging work demonstrated posterior cerebellar activation ^5,6^ during a number of non-motor tasks, such as working memory ^5^, inhibition ^7^, and shifting ^8^. Further, the cerebellum is both functionally and structurally connected with the cortex in a topographically consistent manner ^9^. Critically, closed loop cerebello-thalamo-cortical circuits between cortical regions involved in cognition (i.e. prefrontal cortex) and the cerebellum have been found ^10–12^, providing the underlying connections for cerebello-cortical interactions during task performance.

The cerebello-thalamo-cortical circuit is purported to allow for the processing of internal models created in the cerebellum ^10,13^, which allow for smooth movement and, based on our current understanding, efficient cognition as well. Briefly, it is believed that the cerebellum encodes internal models (i.e. forward or inverse) of processes completed by the cortex ^13^, such that a copy of the information used by the cortex is sent and stored in the cerebellum ^10^, in order to complete a process precisely. A forward model will use these copies to predict the expected result of a motor command but uses error learning to update the command if the predicted outcome does not match the desired outcome. An inverse model will choose the copies necessary to achieve a desired outcome. Critically, these internal models are continually updated based on an input-output relationship between motor commands and their consequences ^13^. Any degradation of these models may contribute to behavioral decline and performance decrements (Bernard et al., 2020; Bernard & Seidler, 2014; Ilg et al., 2013; Schmahmann & Sherman, 1998). This theory of internal models is broadly used to account for the role the cerebellum plays in non-motor cognitive processing, as the cerebellum might be involved in functionally distinct computations ^16^.

Despite a growing literature demonstrating cerebellar activation during nonmotor cognitive processing, little work has investigated how the cerebellum relates to cortical processing. Past work suggests the cerebellum might provide the cortex with processing resources, such that when output from the cerebellum is degraded, performance suffers ^14,15,17^. That is, when tasks become more automatic, individuals can rely more on internal models and cerebellar processing, freeing up cortical resources, particularly if tasks become increasingly complicated. Not only do internal models allow for smooth, efficient, and automatic performance, but they also free up more cortical resources for other processing. Recent advancements in non-invasive stimulation, such as transcranial direct current stimulation (tDCS) allow us to further explore the role of the cerebellum in cognition. tDCS increases (anodal) or decreases (cathodal) neural activity in the cerebral cortex using a small amount of electrical current, which in turn can impact behavior (Coffman et al., 2014). However, the cellular structure of the cerebellum seems to reverse this effect. Specifically, optogenetic work in rodents demonstrates that anodal stimulation excites inhibitory cells in the cerebellum which in turn results in decreased signal to the cortex. Conversely, cathodal stimulation inhibits the inhibitory cells, resulting in increased signal to the cortex ^19^.

Behavioral effects in the human brain broadly fall in line with the polarity specific effects of cerebellar tDCS (Ballard et al., 2019; Cantarero et al., 2015; Ferrucci et al., 2013; Pope & Miall, 2012; Shah et al., 2013). During sequence learning, cathodal stimulation results in task improvement, while anodal stimulation hampers performance ^23^, particularly during late learning ^21^. Further, work examining non-motor cognitive processing has also demonstrated that cathodal stimulation over the right cerebellum improves performance, and anodal stimulation impairs performance (Ferrucci et al., 2008; Pope & Miall, 2012). While there are clear behavioral effects, it is not clear what neural processing occurs to give rise to these behavioral effects.

Imaging work in conjunction with tDCS is limited ^26–30^. When combining tDCS with fMRI, Macher and colleagues found decreased activation in the right cerebellar lobule VIIb during the late encoding phase of a phonological storage task after anodal stimulation ^29^. Cathodal stimulation leads to disinhibition of the dentate nucleus ^30^, in line with past optogenetic work ^19,31^. Though limited, cerebellar stimulation resulted in activation changes, though to this point, the results have been limited to the cerebellum itself, and impacts on cortical activation patterns remain unclear.

Here, we are interested in understanding how the cerebellum interacts with the cortex to support cortical processing. We are testing the idea that the cerebellum supports cortical processing, such that when the cerebellum is functioning properly, it can free up cortical resources and help maintain task performance. With this idea in mind, we would thus expect that in addition to impacts on behavior, downregulation of the cerebellum may result in the need for additional cortical resources to perform a task. To investigate this, we combined cerebellar tDCS and fMRI. In the current study, participants were placed in one of three stimulation conditions (anodal, cathodal, or sham) and completed both motor (sequence learning) and non-motor (Sternberg) tasks to better understand how the availability of cerebellar processing resources impact cortical processing. Stimulation was applied to the right cerebellum. In line with previous optogenetic work ^19,31^, we predict increased cortical activation following cathodal stimulation to the right cerebellum and decreased activation following anodal stimulation during the performance of both the sequence learning and Sternberg tasks.

## Methods

### Participants

Seventy-five healthy, young adults participated in this study and were provided monetary compensation for their time. Exclusion criteria included left handedness, history of neurological or mood disorders, skin conditions, and history of concussion. Data was not collected for one participant because the participant did not wish to complete the experiment after providing consent. For the Sternberg data, an additional two participants were not analyzed because task accuracy was below 20% (n=1) and a computer error interrupted data recording (n=1). Three participants were excluded from the sequence learning analysis due to computer errors (n=2) and excessive movement (n=1). Thus, seventy-four right-handed participants (38 female) ages 18 to 30 (*M*= 22.03 years, *SD*= 3.44) were included in the analyses. Participants were randomly assigned to either the anodal (sequence n=25; Sternberg n=23), cathodal (sequence n=24; Sternberg n=25), or sham (sequence n=22; Sternberg n=24) stimulation condition. All procedures completed by participants were approved by the Texas A&M University Institutional Review Board and conducted according to the principles expressed in the Declaration of Helsinki.

### Procedure

The entire experiment took approximately two hours to complete. Stimulation was completed and behavioral data were collected within 80 minutes. Following the completion of the consent form, participants completed a basic demographic survey, followed by tDCS (see below for details). Participants were blind to the stimulation types. After stimulation, participants completed a computerized Sternberg ^32^ and sequence learning ^33^ task in the MRI environment while brain imaging data were collected. Tasks were administered in a pre-determined random order (for more details, see below).

### tDCS Stimulation Parameters

Cathodal, anodal, or sham stimulation was administered using a Soterix 1×1 tES system. Each electrode was placed in a saline soaked sponge (6 mL per side), with the stimulation electrode placed two cm below and four cm lateral of the inion over the right cerebellum, and the return electrode placed on the right deltoid (Ferrucci, Cortese, & Priori, 2015).

To ensure proper connection with the scalp, an initial 1.0 mA current was set for 30 seconds. If contact quality is below 40%, adjustments, such as moving hair to increase the electrode’s contact with the scalp, were made and contact quality was rechecked. Following a successful re-check, participants completed a 20-minute stimulation session at 2 mA (Ferrucci et al., 2015; Grimaldi et al., 2014, 2016). During the stimulation conditions, maximum stimulation intensity was reached in 30 seconds and maintained for 20 minutes, and then would return to 0 mA. During the sham condition, maximum stimulation intensity would be reached, but would then immediately return to 0 mA. There was no additional stimulation during the 20-minute session. Stimulation was followed by the completion of the behavioral tasks in the scanner.

### Behavioral Tasks

Task administration started about 20 minutes after stimulation and the tasks took approximately 35 minutes to complete. This is within the 90 minute window in which stimulation is thought to be effective ^36^. However, task order was counterbalanced across participants to mitigate the impact of time after stimulation on task performance.

#### Sequence Learning

The sequence learning task ^33^ was administered via computer using PsychoPy (Peirce et al., 2019; Peirce, 2007). Participants were shown four empty rectangles and instructed to indicate the location of the rectangle that was filled as quickly as possible via button press. Though the stimuli were presented for 200ms, the participant had 800ms to respond before the next stimulus appeared. Random blocks (R) had 18 trials and sequence (S) blocks had 36 trials. During sequence trials, participants had to learn a six-element sequence (1-3-2-3-4-2), which was repeated six times within a block. The order of the task was as follows: R-S-S-S-R-R-S-S-S-R-R-S-S-S-R. For the purposes of analysis here, the first three sequence blocks were considered early learning, the central sequence blocks were middle learning, and the last sequence blocks were considered late learning ^39^. Briefly, early learning is marked by more cognitively focused activities that necessitate active thinking and working memory ^40–42^. As the skill becomes automatic via repetition and practice, the late learning phase becomes increasingly motor-focused. Dependent variables used to estimate learning were mean reaction time for correct trials and average total accuracy.

#### Sternberg

The Sternberg Task ^32^ was administered via computer using PsychoPy v3.1.2 (Peirce et al., 2019; Peirce, 2007). At the beginning of a trial, participants were given six seconds to remember a string of either one, five or seven capitalized letters, which represent low, medium, and high load, respectively. Following the presentation of the study letters, participants were shown individual lower-case letters and told to indicate whether the letter was one of the study letters shown at the beginning of the trial, via button press. Each letter was displayed for 1200ms, separated by a fixation cross that lasted 800ms. Each participant completed three runs of this task. Within each run, a participant completed three blocks of 25 trials each, for a total of 225 trials. Dependent variables were average reaction time for correct trials and accuracy.

#### Behavioral Data Analysis

Statistical analyses were conducted in R ^43^, using the *lme4* ^44^ package, and p-value estimates were determined using the *lmerTest* package ^45^. A *p*<.05 threshold was used as the cut-off for significance. When necessary, the *emmeans* package ^46^ was used to follow-up on significant effects. These comparisons of estimated marginal means used Bonferroni-corrected *p*-values.

Task data was analyzed using liner mixed effects models using restricted maximum likelihood, as it produces unbiased estimates of variance and covariance parameters, ideal for mixed effect models with small samples. Learning phase (early, middle, late) was included as a fixed factor for the sequence task, with all random trials included for comparison. Load (low, medium, and high) was included as a fixed factor for the Sternberg task. Stimulation type (cathodal, anodal, or sham stimulation) was included as a fixed effect and subject was included as a random effect for both tasks. A model was completed for both reaction time for correct trials and accuracy across both tasks.

### fMRI Procedures

#### Data Acquisition

fMRI data was collected at the Texas A&M Translational Imaging Center with a 3-T Siemens Magnetom Verio scanner using a 32-channel head coil. Three scans with alternate phase encoding directions were used to collect blood oxygen level dependent (BOLD) whole brain scans with a multiband factor of 4 (number of volumes = 134, repetition time [TR] = 2000 ms, echo time [TE] = 27 ms; flip angle [FA] = 52°, 3.0 × 3.0 × 3.0 mm3 voxels; 56 slices, interleaved, slice thickness=3.00mm, field of view (FOV) = 300 × 300 mm; time = 4:40 min). An additional high resolution T1-weighted whole brain anatomical scan was taken (sagittal; GRAPPA with acceleration factor of 2; TR = 2400 ms; TE = 2.07 ms; 0.8 × 0.8 × 0.8 mm3 voxels; 208 slices, interleaved, slice thickness= 0.8; FOV = 256 × 256 mm; FA = 8°; time = 7:02 min) for data normalization.

#### fMRI data pre-processing and analysis

Images were converted from DICOM format to NIFTI files and organized into a Brain Imaging Data Structure (BIDS) using bidskit (v 2019.8.16; Mike Tyszak, 2016). Functional images were encoded using opposite phase encoding directions. For distortion correction, single 4D images were taken for each participant from each phase encoding direction and were merged. Then fieldmap images were created using FSL’s topup to unwrap images (Andersson et al., 2003).

FMRI data was processed using FEAT (FMRI Expert Analysis Tool) Version 6.00, part of FSL (FMRIB’s Software Library, www.fmrib.ox.ac.uk/fsl). Registration to high resolution structural and/or standard space images was carried out using FLIRT ^48,49^. Registration from high resolution structural to standard space was then further refined using FNIRT nonlinear registration (Andersson, Jenkinson, & Smith, 2007; Andersson, Jenkinson, & Smith, 2007). The following pre-statistics processing was applied: motion correction using MCFLIRT ^49^; slice-timing correction using Fourier-space time-series phase-shifting; non-brain removal using BET ^52^; spatial smoothing using a Gaussian kernel of FWHM 5mm; grand-mean intensity normalization of the entire 4D dataset by a single multiplicative factor. ICA was carried out using MELODIC ^53^, to investigate the possible presence of unexpected artefacts or activation. Time-series statistical analysis was carried out using FILM with local autocorrelation correction ^54^. Subject level variables were modeled using fix effects and group level comparisons were modeled using FLAME 1 & 2 mixed effects. The subject level contrast collapsed across stimulation condition and contrasted activation conditions between sequence learning phases types (early, middle and late) and working memory loads (low, medium, high). Group level analyses (cathodal, anodal, sham) were thresholded non-parametrically using clusters determined by *z*>3.1 and a (corrected) cluster significance threshold of *p*=0.05 ^55^. For display purposes, cortical volumetric maps were projected on the HCP 1200 Subject Group Average Pial Surface ^56^ using the Connectome Workbench v. 1.5.0 (https://www.humanconnectome.org/software/get-connectomeworkbench). The 2-D cerebellar slices were created using MRICron (https://www.nitrc.org/projects/mricron) on the ch2bet template ^57^.

#### ROI Analysis

Region of interest (ROI) analyses were also conducted to determine if stimulation affected signal in specific ROIs. Here, we used masks that covered bilateral parietal cortices, bilateral frontal cortices, and bilateral Crus I using masks from an existing repository of functional ROIs (Figure 1; Shirer et al., 2012), provided by the FIND Lab (findlab.stanford.edu). The mask used for the frontal lobes cover the left and right Middle Frontal Gyrus, and the left and right Superior Frontal Gyrus (Broadman’s area 8, 9, 46). Similarly, the masks used for the parietal lobes cover the left and right Inferior Parietal Gyrus, right Supramarginal Gyrus, left Inferior Parietal Gyrus, left Precuneus, and left and right Angular Gyrus (Broadman’s area 7, 39, 40). The masks used for the frontal and parietal lobe analyses were pulled from the Executive Control Network repository. Cerebellar masks were pulled from the Language repository (left Crus I) and the left Executive Control Network repository (right Crus I).

**Figure 1.**
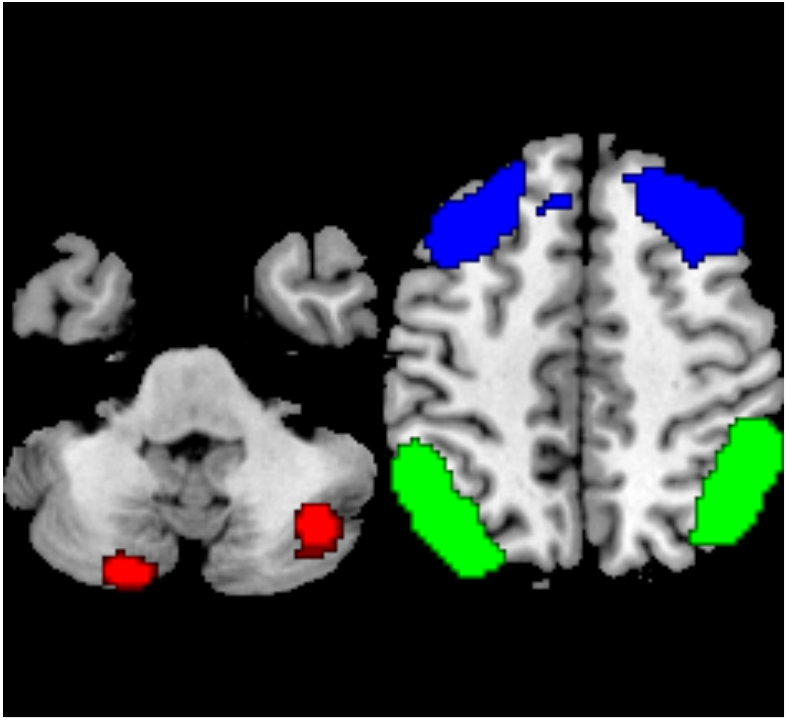
Regions of interest used to examine percent signal change (Shirer et al., 2012). Red= Crus I; Blue=Frontal Gyrus; Green=Parietal Gyrus.

These masks were fed into FSL’s Featquery, which calculated percent signal change for each subject for both the sequence learning and Sternberg task. ANOVAs were conducted to determine if there were significant differences in mean percent signal change in an ROI within each stimulation condition (cathodal, anodal, sham). Mean percent signal change was assessed for each individual (i.e. left Crus I) and combined (i.e. left and right frontal lobe) ROI, in order to look at signal change within hemispheres (i.e. left Crus I) and possible interactions between hemispheres (i.e. left and right frontal lobe). Pearson correlations were also run for each ROI within each stimulation condition to see if performance was related to signal change. These comparisons were corrected using Bonferroni-corrected *p*-values.

## Results

One issue that might affect behavioral outcomes is task order and stimulation decay, given the time between completion of the tDCS session and the two tasks in the scanner. Even though task order was counterbalanced to mitigate any effect of stimulation decay, it is still possible to that the effect of stimulation was lessened during the second task completed in the scanner. We investigated this to quantify the impacts of stimulation on task performance over time. Please see the Supplemental Materials for details. In brief, the impact of task order was minimal, at best, and highly limited, suggesting that stimulation effects persisted over the course of the scanning session.

### Sequence Learning

#### Behavioral

Mean reaction times (RT) and accuracy for the sequence learning task can be found in Supplemental Table 1 and in Supplemental Figure 3. First, we found a significant phase by stimulation interaction (*F*(6, 28,332) = 4.96, *p* < .001), such that the magnitude of change in RT is significantly greater following cathodal stimulation between middle learning and random button presses, compared to anodal and sham. Additionally we found a significant effect of learning phase (*F*(3, 28,332) = 1,796.79, *p* < .001), such that reaction times for early, middle, late, and random learning trials were all significantly different from one another (*p*s < .001). There was no effect of stimulation (*F*(2, 68) = 0.693, *p* = .504).

When examining accuracy, we found a phase by stimulation interaction (*F*(6, 30,592) = 3.74, *p* = .001), such that accuracy was lower during late learning following anodal stimulation, compared to sham (*p* =.020) and cathodal (*p* < .002) stimulation. There was no main effect of stimulation on accuracy (*F*(3, 68) = 1.54, *p* = .223), though we did find an effect of phase (*F*(3, 30,592) = 16.15, *p* < .001), such that accuracy was worse for early learning, followed by late learning, and lastly middle learning. Taken together, RT was improved following cathodal stimulation during middle learning and accuracy was negatively impacted following anodal stimulation during later learning phases. More generally, accuracy data also suggest that participants learned across the course of the task.

#### Imaging

Patterns of brain activation after stimulation for the three learning phases are presented in the Supplementary material. In brief, activation patterns were consistent with canonical findings of cortical motor activation ^59,60^, and patterns typically seen during explicit motor sequence learning ^61^ for both sham and active stimulation conditions. Here, we have focused on patterns of activation comparing sequence to random blocks after stimulation. Please see Table 1 for detailed reporting of activation foci and statistics.

**Table 1.** Significant contrast clusters following stimulation during a Sequence learning task.

| Phase | Stimulation | Region | BA | Voxels | MNI Coordinates |  |  | Z |
| --- | --- | --- | --- | --- | --- | --- | --- | --- |
|  |  |  |  |  | x | y | z |  |
| Sequence<br>> Random | Cathodal | Left Thalamus | Left-Thalamus (50) | 259 | -6 | -26 | 10 | 4.93 |
|  |  | Left Supplementary motor area | Right-BA6 | 153 | 0 | -6 | 56 | 5.26 |
|  | Anodal | Left Inferior parietal, but<br>supramarginal and angular gyri | Left-BA40 | 1987 | -40 | -40 | 42 | 7.13 |
|  |  | Left Supplementary motor area | Left-BA8 | 932 | -6 | 16 | 44 | 6.11 |
|  |  | Right Inferior parietal, but<br>supramarginal and angular gyri | Right-BA7 | 819 | 36 | -46 | 42 | 6.93 |
|  |  | Left Inferior occipital gyrus | Left-BA19 | 464 | -46 | -72 | -4 | 6.26 |
|  |  | Right Fusiform gyrus | Right-Fusiform (37) | 329 | 38 | -70 | -16 | 5.63 |
|  |  | Left Insula | Left-BA45 | 268 | -34 | 18 | 6 | 5.7 |
|  |  | Left Thalamus | Left-Thalamus (50) | 219 | -8 | -16 | 10 | 5.51 |
|  |  | Right Angular gyrus | Right-BA39 | 189 | 36 | -56 | 54 | 5.8 |
|  |  | Right Insula | Right-BA44 | 147 | 40 | 18 | 4 | 5.09 |
|  |  | Right Thalamus | Right-Thalamus (50) | 110 | 10 | -18 | 4 | 5.12 |
|  |  | Left Middle frontal gyrus | Left-BA10 | 100 | -32 | 46 | 24 | 4.84 |
|  | Cathodal<br>> Sham | Right Caudate nucleus | Right-Caudate (48) | 145 | 8 | 0 | 12 | 5.93 |
|  | Anodal ><br>Sham | Right Middle frontal gyrus | Right-BA10 | 116 | 28 | 50 | 2 | 4.83 |
|  |  | Right Lobule VI, cerebellum | Right-Fusiform (37) | 104 | 38 | -44 | -28 | 4.97 |
|  | Anodal ><br>Cathodal | Left Inferior parietal, but<br>supramarginal and angular gyri | Left-BA40 | 446 | -40 | -40 | 42 | 5.67 |
|  |  | Right Angular gyrus | Right-BA39 | 326 | 42 | -48 | 38 | 5.23 |
|  |  | Right Angular gyrus | Right-BA7 | 193 | 28 | -58 | 52 | 4.76 |
|  |  | Left Lobule IV & V, cerebellum | Left-Fusiform (37) | 154 | -20 | -40 | -30 | 4.47 |
|  |  | Left Precentral gyrus | Left-BA6 | 122 | -60 | 6 | 32 | 5.08 |
|  |  | Right Lobule VI, cerebellum | Right-Fusiform (37) | 108 | 32 | -46 | -26 | 5.01 |

##### Sequence>random

When comparing sequence to random blocks (Figure 2a), we found greater bilateral cortical activation following anodal stimulation, with activations in the left middle frontal gyrus, left supplemental motor area, left inferior occipital lobe, right angular, right fusiform gyrus, left and right inferior parietal, and left and right insula. There was also activation in subcortical regions, particularly the thalamus. Following cathodal stimulation (Figure 2b), we found only activations in the left thalamus and left supplemental motor region.

**Figure 2.**
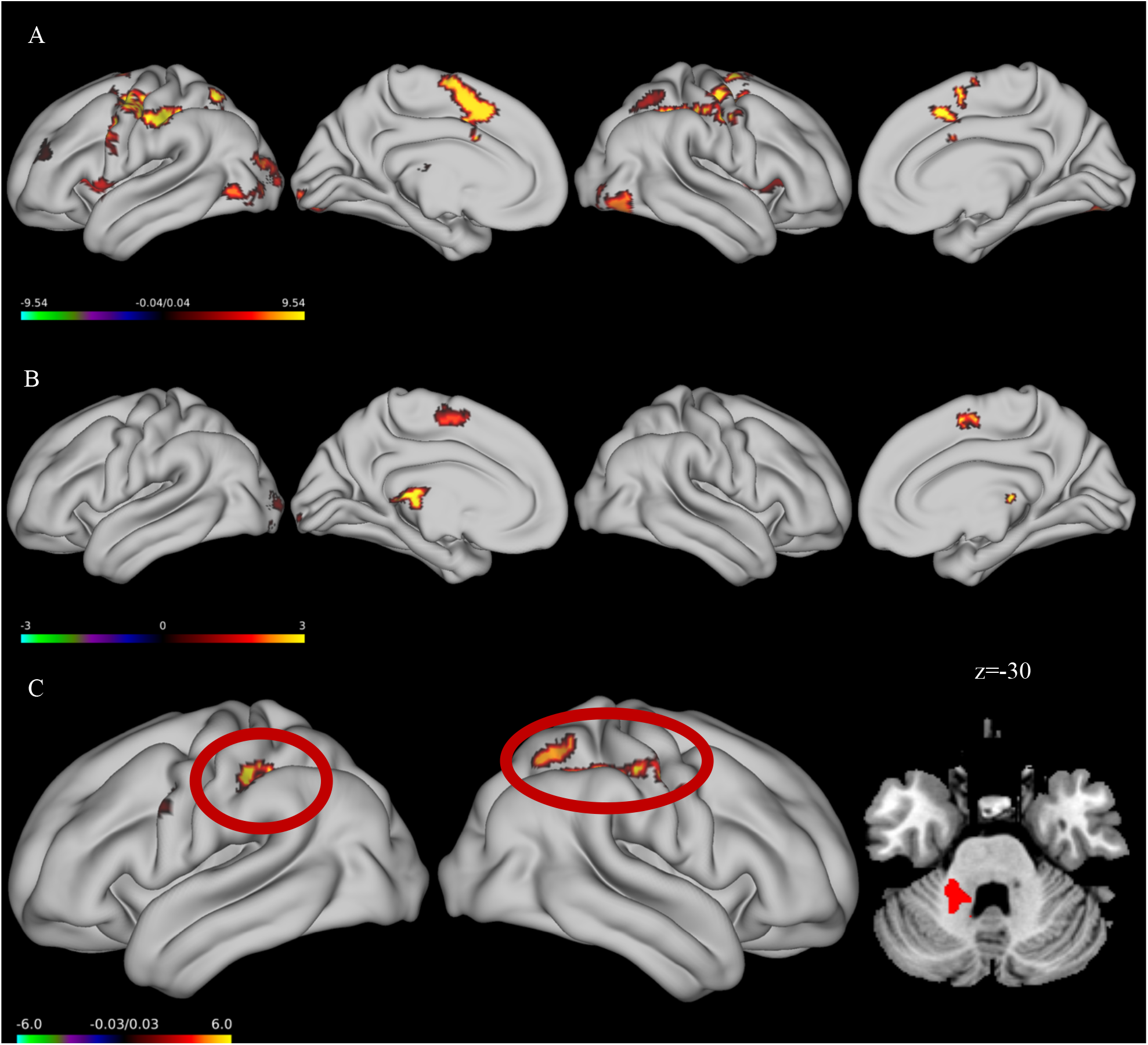
Significant activations greater during sequence learning than random trials (sequence>random); (A) Anodal Stimulation; (B) Cathodal Stimulation; (C) Activations greater following anodal stimulation compared to cathodal stimulation.

Further, we investigated the statistical differences between the stimulation groups (Figure 2c). The anodal stimulation group had greater activation in the right angular gyrus, left inferior parietal lobe, left precentral gyrus, and lobules IV-VI in the cerebellum compared to cathodal stimulation. Further, anodal stimulation resulted in larger activation in the right middle frontal lobe and right lobule VI in the cerebellum when compared to the sham group. This is again consistent with our scaffolding hypothesis, wherein there was additional cortical activation (here, bilaterally in the parietal lobes) after stimulation to the cerebellum that is thought to downregulate its function and output ^19,62^.

### Sternberg Task

#### Behavioral

Mean reaction times and accuracy on the Sternberg task can be found in Supplemental Table 4 and are depicted visually in Supplemental Figure 8. When examining the fixed effects of reaction time, there was a significant effect of load (*F*(2, 9800) = 665.74, *p* < .001), such that reaction times for each load condition were significantly different from each other (*p*s< .001). This demonstrates the increase in difficulty associated with increased load. We also found a significant effect of stimulation (*F*(2, 68.9) = 4.20, *p* = .011), such that reaction times following anodal (*p* = .038) and cathodal (*p* = 0.035) stimulation were both significantly quicker relative to sham. We did not find a stimulation by load interaction (*F*(4, 9800) = 1.05, *p* = .380).

With respect to accuracy, we only found an effect of load (*F*(2, 10,290) = 194.93, *p* < .001), such that accuracy was best on low (*p* < 0.001) load, then medium load (*p* < .001), and then high load (*p* < .001). There was no effect of stimulation (*F*(2, 69) = 0.026, *p* = .974), or a load by stimulation interaction (*F*(4, 10,290) = 1.29, *p* = .271).

#### Imaging

General patterns of activation after stimulation for each load condition is reported in the Supplementary material. Briefly, during sham stimulation trials we found activation in the frontal and parietal regions one would expect when completing a verbal working memory task ^63^. Contrasts between the load conditions in the sham group are also reported in the Supplement. Notably, we demonstrated the expected effects of load wherein activation was significantly higher for high relative to low load blocks. Here, we focused on the impact of stimulation when looking at the contrasts between load levels. Activations are reported in Table 2.

**Table 2.**
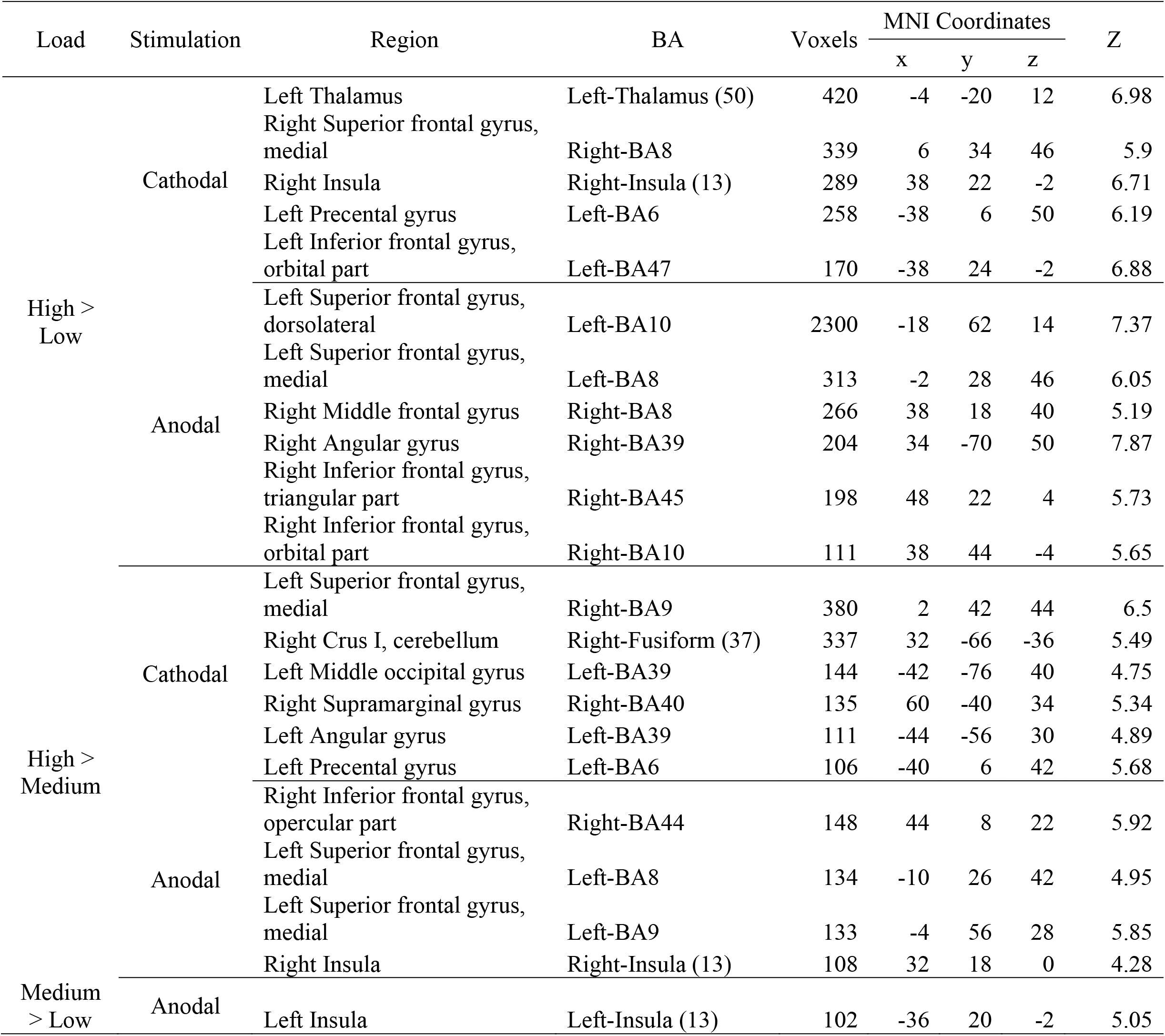
Significant contrast clusters following stimulation during a Sternberg task.

##### High > Low

The anodal group (Figure 3a) showed higher activations under high load in the right angular gyrus. Additionally, there were greater activations in the left superior, left and right inferior, and right middle frontal lobes during high load relative to low in the anodal stimulation group. After cathodal stimulation (Figure 3b) there was greater activation on high load relative to low load blocks in left inferior and right superior frontal regions, left precentral gyrus, left thalamus, and right insula.

**Figure 3.**
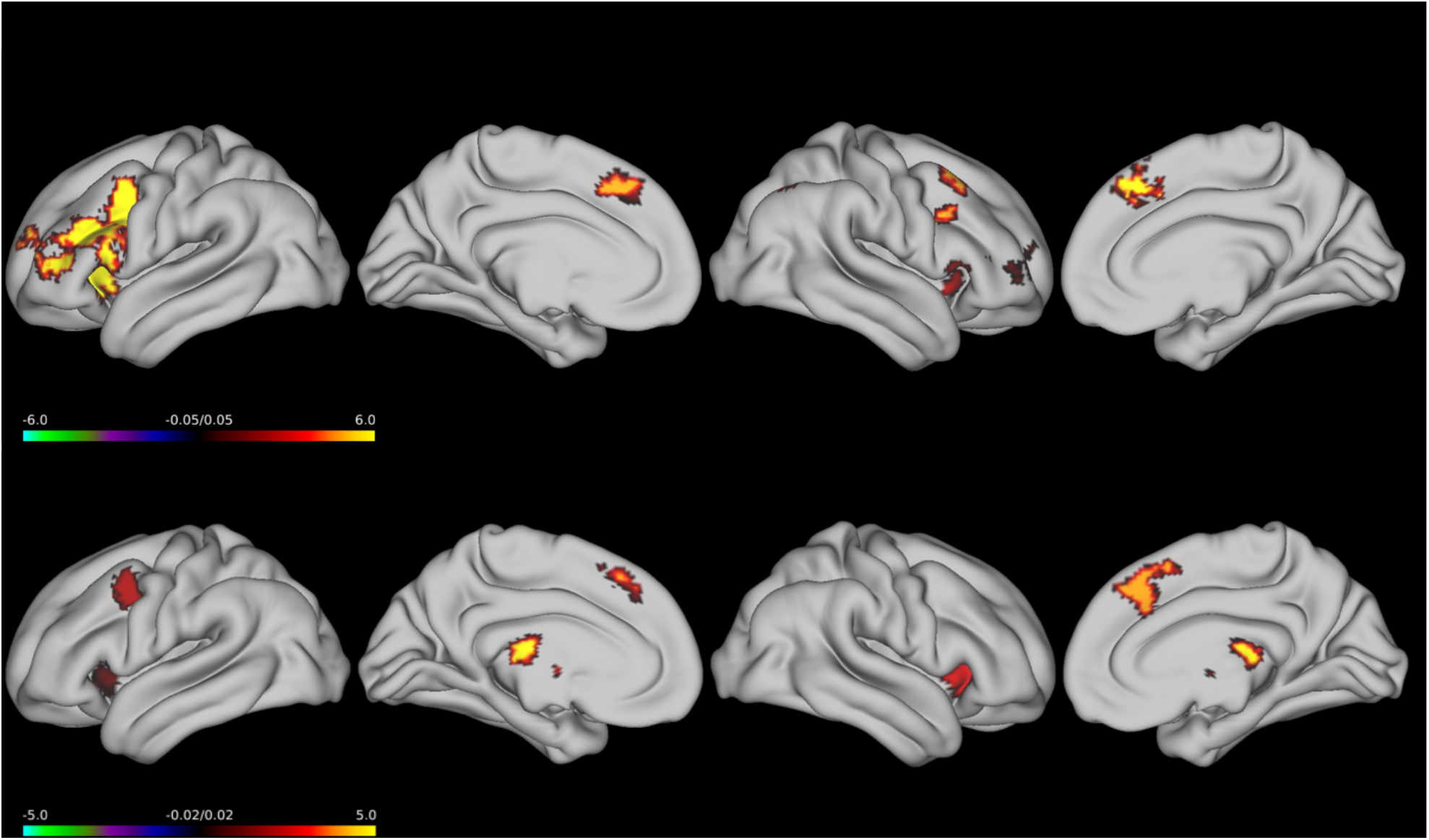
Significant contrast activations for the High > Low load contrast a Sternberg task; (A) Anodal; (B) Cathodal.

##### High > Medium

Following anodal stimulation (Figure 4a), there was greater activation in frontal regions including the right inferior frontal lobe, and the left superior frontal lobe during the high load condition as compared to medium. When comparing high to medium load in the cathodal group (Figure 4b), activation was higher in frontal, occipital, and parietal regions including the left superior frontal lobe, left middle occipital lobe, supramarginal gyrus, left angular, and left precentral lobe during high load compared to medium load. Additionally, greater activation was also seen in the right Crus I in the cerebellum.

**Figure 4.**
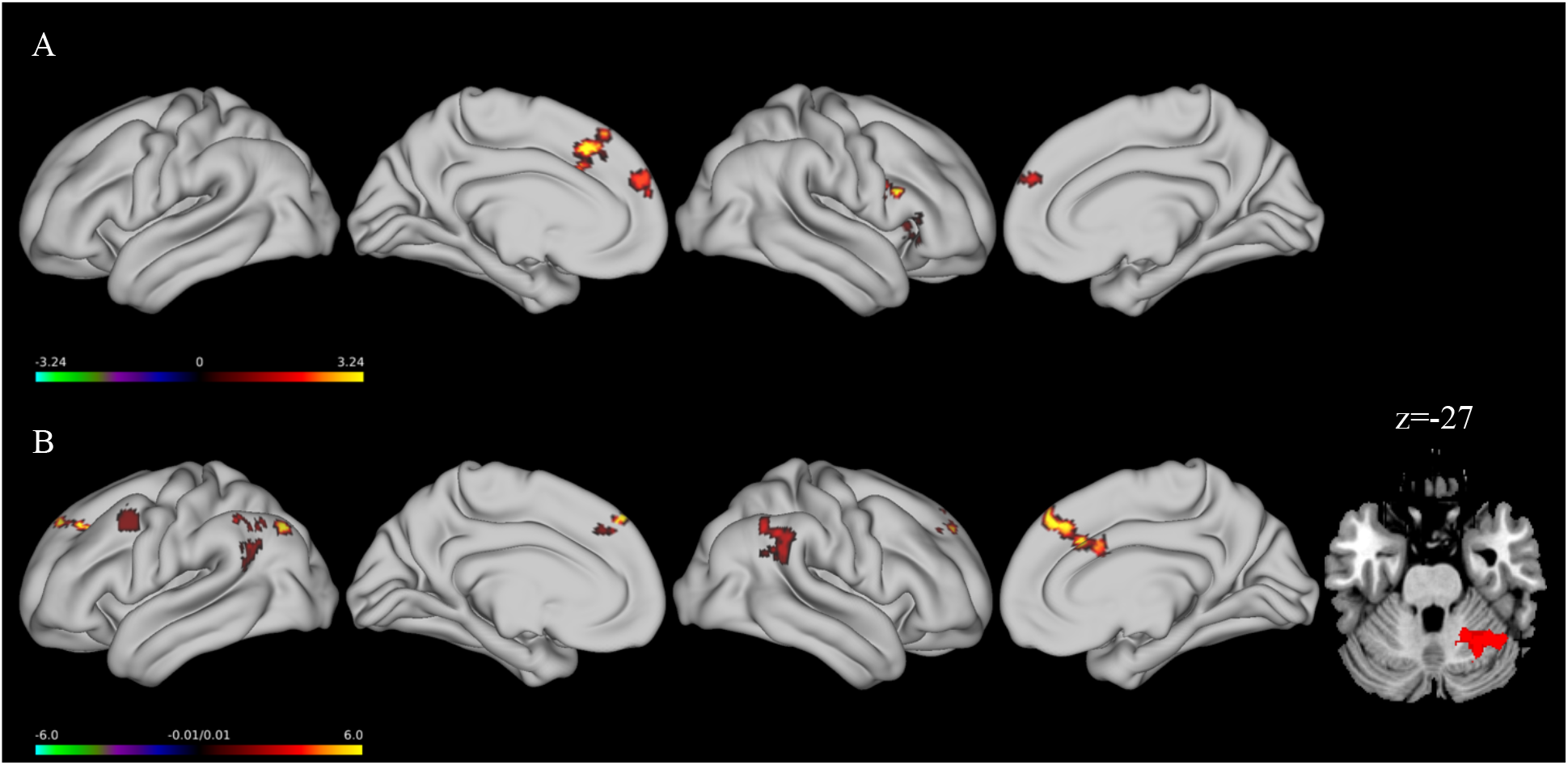
Significant contrast activations for the High > Medium load contrast a Sternberg task; (A) Anodal; (B) Cathodal

##### Medium > Low

Here, the anodal stimulation group showed greater activation in the left insula under medium load compared to low load. There were no significant activation differences following cathodal stimulation.

Together, though there were no behavioral effects of stimulation, frontal and parietal activations were greater when processing was high, and in the anodal stimulation group, we saw bilateral activation of the frontal lobes (Figure 3a), consistent with our scaffolding hypothesis wherein additional resources may be needed to make up for the down regulation of the cerebellum with anodal tDCS ^17,62^.

#### ROI analysis

A correlation analysis was also conducted to see if signal change was associated with Sternberg or sequence learning performance (Table 3). During Sternberg task performance, accuracy increased as signal increased in the left parietal lobe following cathodal stimulation (*p* =.013; Figure 5a), but decreased as signal increased following anodal stimulation (*p* =.050). During sequence learning, reaction time increased as signal increased in the right parietal lobe (*p* =.014; Figure 5b) following anodal stimulation. However, these effects did not survive a Bonferroni multiple comparison correction.

**Table 3.** Correlation matrix for percent signal change in Crus I, frontal and parietal regions during task performance. P-values displayed are uncorrected.

| **Region** | **Hemisphere** | **Load** | **Stimulation** | **DV** | **r** | **p** | **Task** |
| --- | --- | --- | --- | --- | --- | --- | --- |
| Crus I | Left | Low | Cathodal | Accuracy | 0.477 | 0.016 | Sternberg |
| Crus I | Right | High | Sham | Reaction Time | 0.529 | 0.008 | Sternberg |
| Crus I | Right | Medium | Cathodal | Accuracy | 0.396 | 0.05 | Sternberg |
| Frontal | Left | Low | Anodal | Reaction Time | -0.423 | 0.044 | Sternberg |
| Frontal | Right | High | Anodal | Accuracy | -0.482 | 0.02 | Sternberg |
| Frontal | Right | Low | Cathodal | Accuracy | 0.421 | 0.036 | Sternberg |
| Parietal | Left | All | Anodal | Accuracy | -0.413 | 0.05 | Sternberg |
| Parietal | Left | All | Cathodal | Accuracy | 0.491 | 0.013 | Sternberg |
| Parietal | Left | High | Anodal | Accuracy | -0.52 | 0.011 | Sternberg |
| Parietal | Left | Medium | Anodal | Reaction Time | 0.429 | 0.041 | Sternberg |
| Parietal | Left | Medium | Cathodal | Accuracy | 0.468 | 0.018 | Sternberg |
| Parietal | Right | High | Anodal | Accuracy | -0.576 | 0.004 | Sternberg |
| Parietal | Right | Low | Cathodal | Accuracy | 0.404 | 0.045 | Sternberg |
| **Region** | **Hemisphere** | **Load** | **Stimulation** | **DV** | **r** | **p** | **Task** |
| Frontal | Left | Early | Sham | Accuracy | -0.443 | 0.039 | Sequence |
| Frontal | Right | Early | Cathodal | Accuracy | 0.628 | 0.001 | Sequence |
| Parietal | Left | Late | Anodal | Reaction Time | 0.533 | 0.006 | Sequence |
| Parietal | Left | Middle | Sham | Accuracy | -0.497 | 0.019 | Sequence |
| Parietal | Right | All | Anodal | Reaction Time | 0.483 | 0.014 | Sequence |
| Parietal | Right | Late | Anodal | Reaction Time | 0.596 | 0.002 | Sequence |
| Parietal | Right | Middle | Anodal | Reaction Time | 0.489 | 0.013 | Sequence |
| Parietal | Right | All | Sham | Accuracy | -0.511 | 0.015 | Sequence |
| Parietal | Right | Early | Sham | Accuracy | -0.453 | 0.034 | Sequence |

**Figure 5.**
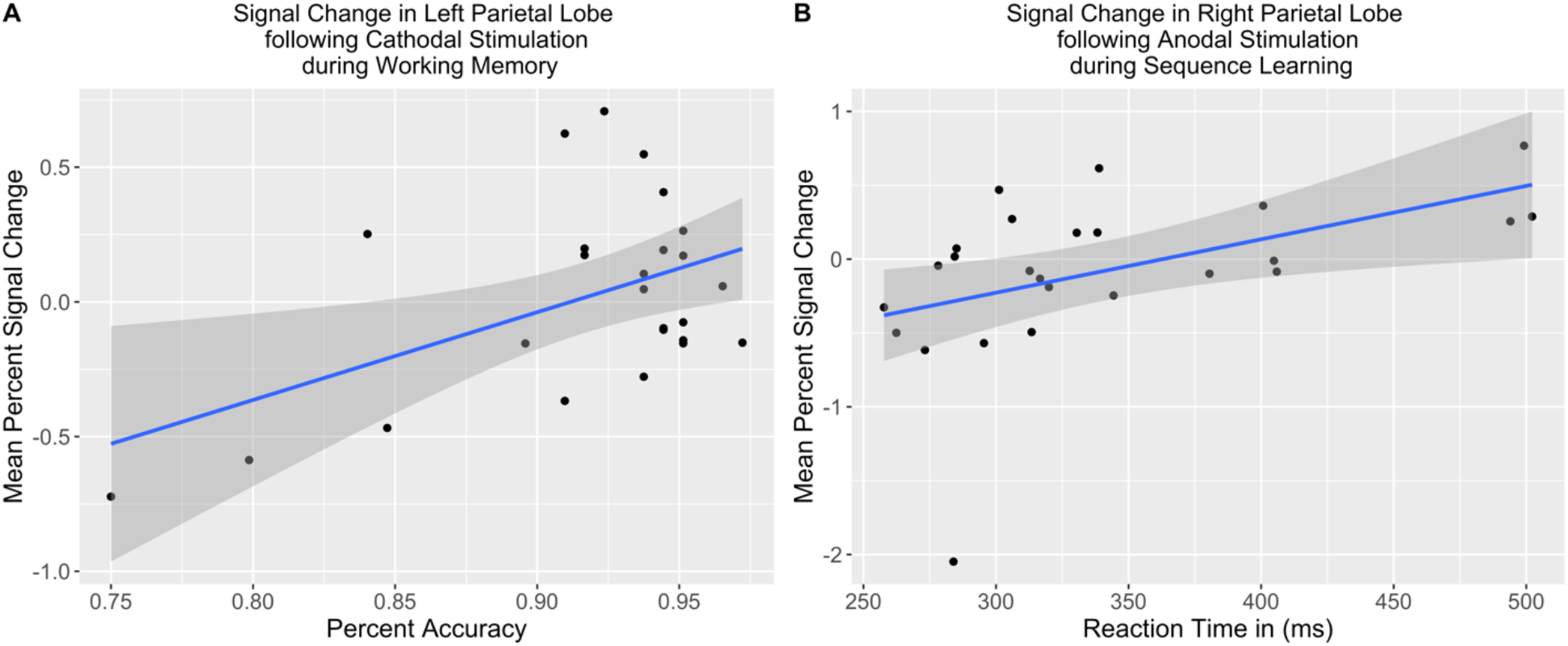
Correlations between mean percent signal change and task performance following stimulation. (A) Mean percent signal change in the left parietal lobe following cathodal stimulation during a working memory task. (B) Mean percent signal change in the right parietal lobe following anodal stimulation during a sequence learning task.

## Discussion

The literature implicating the cerebellum in cognitive processing is growing (Buckner, 2013; Schmahmann et al., 2019; Stoodley et al., 2012b), but little work has examined how activation in this structure relates to that in the cortex during non-motor tasks. Recent aging work has implicated the cerebellum in cortical scaffolding ^14,15,17^, suggesting that the cerebellum is recruited as a support system for processing through the use of internal models and more automatized processing. Here, we combined tDCS and fMRI to better understand how activation patterns might relate to behavioral performance and to understand what role the cerebellum might play in cognitive processing, particularly in conjunction with processing in the cerebral cortex. Following anodal, cathodal, or sham stimulation, participants completed a motor learning (explicit sequence learning) or verbal working memory (Sternberg) task. Broadly, we found increased cortical activation following anodal stimulation (thought to downregulate cerebellar function) across task domains, implicating the cerebellum as a critical scaffold for cortical processing ^14,17^, particularly when cerebellar output is thought to be degraded. Together, this work provides novel insights into the potential cerebellar scaffolding mechanism. Results and implications are discussed below.

### Working Memory

Behaviorally, we found the expected effect of load, such that performance (both reaction time and accuracy) was best for low load, followed by medium load, and finally worst for high load. We did not find an effect of stimulation on accuracy during performance of the Sternberg task, but we did find that both anodal and cathodal stimulation improved reaction time. Though we acknowledge that reaction time is not the only measure of working memory performance, the current data mimic the mixed nature of this literature. Regarding accuracy, there was no effect of stimulation, though accuracy was high, perhaps making it difficult for stimulation to modulate task performance.

Activations following stimulation (both cathodal and anodal) are consistent with past work investigating working memory ^63,65^ and could explain the behavioral effect we found on reaction time. Functional patterns following cathodal stimulation under high load, showed activations in frontal, parietal and cerebellar regions associated with verbal working memory ^65^. However, when contrasting high and low load following anodal stimulation, we found greater frontal activations to the inferior, middle and frontal lobes, which are also regions implicated in verbal working memory task performance ^63,65^. We propose that this increased bilateral cortical activation may be compensation as a result of diminished cerebellar output, given what is known about the impact of anodal stimulation on the cerebellum ^19,31^. In the current work, this compensatory response following anodal stimulation might have been effective and helped improve behavioural performance as measured by reaction time. Cathodal stimulation activated expected regions, but also in turn by stimulating cerebellar activity, may have positively influenced motor performance. Work by Macher and colleagues applied anodal stimulation to the right cerebellum which resulted in poorer performance on a modified Sternberg task ^29^. Critically, this work also found attenuated signal in the right cerebellum, and decreased functional connectivity to the posterior parietal cortex following anodal stimulation. This attenuated signal to the cortex following anodal stimulation is in line with our predictions. In the current work, it is possible that connectivity to the frontal lobes was also attenuated following anodal stimulation, as might be predicted by Grimaldi et al (2016), resulting in the need for more cortical processing to ensure successful task completion. That is, if the cerebellum was not processing information from the cortex adequately, more cortical resources would be needed to make up for this, resulting in increased cortical activation. Here, cathodal stimulation did result in a behavioral boost manifesting as improved reaction time during a verbal working memory task. But we might also be observing an effective compensatory response in the frontal lobes, such that when anodal stimulation diminished cerebellar output and interrupted processing during high cognitive load tasks, the cortex was able to successfully recruit more neural resources, which ultimately improved reaction time following anodal stimulation.

### Sequence Learning

Behaviorally, we found no significant effect of stimulation on reaction time, but we did find that the anodal stimulation group showed significantly worse accuracy during the late phase of learning. This is consistent with previous findings from our group ^21^, and methodological discrepancies with respect to electrode placement may explain differences from prior work (Ferrucci et al., 2013). In brief, we suggest that anodal stimulation disrupts the formation of internal models during early learning when the cerebellum is particularly active (Imamizu et al., 2000; Ballard et al., 2019), and in late learning when performance would be more automatic, these models cannot be relied upon. Notably, in support of our scaffolding hypothesis, after anodal stimulation, we saw increased frontal and parietal activation, suggesting cortical areas may be compensating for decreased cerebellar processing.

Our imaging demonstrated that anodal stimulation increased cortical activations, in key frontal ^63,65^, parietal ^66^ and cerebellar ^5^ regions associated with non-motor cognition. The activations in parietal regions are greater following anodal stimulation when compared to cathodal, demonstrating how disruptive anodal stimulation might be to task performance and cortical processing. This mimics the effect we see during the Sternberg task, in which anodal stimulation seems to increase cortical activation, presumably due to degraded cerebellar output and associated processing, in an effort to maintain task performance. Perhaps the cerebellum is supporting the working memory processes needed to complete a sequence learning task, but anodal stimulation degrades the cerebellar output necessary to support this process, requiring increased activation in parietal regions associated with working memory ^66^. This provides further evidence to suggest that the cerebellum plays a supporting, scaffolding, role in mon-motor cognitive processing and degradation of cerebellar output has functional and behavioral consequences.

Contrary to the current findings, recent work found anodal stimulation improves sequence learning, particularly in middle to late learning phases ^67^. We should note however, Liebrand and colleagues showed changes in activation that are consistent with increased cortical activation because of degraded cerebellar output following anodal tDCS. Therefore, it is possible that anodal stimulation did modulate cortical activation similar to what is found in the current work, but the behavioral outcomes were negated, due to methodological differences, such as the online nature of stimulation in the work conducted by Liebrand and colleagues ^68^.

Here, we see increased bilateral cortical activation in parietal regions typically active during sequence learning ^66^. Though we saw an increase in activation with anodal stimulation that we argue is compensation for the negative impact on cerebellar processing, the compensation was not enough, which resulted in poor performance in late learning. We speculate that internal models were not adequately created during the earlier phases of learning resulting in a greater need for cortical processing, but it might also explain why accuracy still suffered, as not enough cortical resources were brought on. Since the cerebellum is less able to provide resources to the cortex, cortical activations increase to complete a task efficiently ^17,62^. Critically, we see a decrease in accuracy in late learning following anodal stimulation, which might be a behavioral consequence of degraded cerebellar output, especially when the cortex is not able to fully compensate for the loss of cerebellar resources.

### The cerebellum as a scaffolding structure

The current work looked to better understand the role the cerebellum plays in cognitive processing. In both a verbal working memory and sequence learning task, we have found that anodal stimulation resulted in increased bilateral cortical activation, in regions previously associated with these tasks ^63,65,66^, compared to sham or cathodal stimulation. Optogenetic work suggests that anodal stimulation to the cerebellum will excite inhibitory Purkinje cells, ultimately decreasing signal to the cortex ^19,31^. Our work here suggests that the cortex may compensate for this lost input and processing from the cerebellum by increasing cortical activation. Therefore, it is possible that the cerebellum provides additional resources to help support cortical processing. Specifically, internal models stored in the cerebellum are used for greater automaticity on well learned tasks ^10,13^. However, when cerebellar outputs are degraded, there are negative behavioral implications ^14,17^. Interestingly, some of this increased cortical activation occurred when cognitive processing demands were higher. This was particularly notable during high load in the Sternberg task, perhaps indicating that the cerebellum is increasingly relied upon when cortical regions are taxed. That is, some offloading of processing via internal models may occur when tasks get more difficult. If this is inhibited due to cerebellar dysfunction, age differences, or, as was the case here, due to stimulation, more cortical resources may be needed to maintain performance.

Past work in aging ^62^ and disease ^69^ has suggested that degraded cerebellar output negatively impacts cortical connectivity and activation. Cerebellar resources might be important for cortical processing, as they may provide crucial scaffolding for performance and function ^17,62^. Indeed, a recent imaging meta-analysis indicated that the cerebellum in advanced age might be able to engage in compensatory scaffolding for motor tasks, but there was decreased overlap across cognitive tasks compared to young adults, perhaps contributing to performance differences ^15^. Compensatory scaffolding models argue that deteriorating neural structures in advanced age result in a need for increased activations in cortical regions to compensate for these losses ^70,71^. Based on the current work, anodal tDCS seems to mimic this disrupted cerebellar function, ultimately decreasing cerebellar output, which disrupts cortical processing. This then resulted in the need for increased cortical activation, to maintain task performance. We suggest that anodal stimulation negatively impacts output of the cerebellum via closed-loop circuits with the cortex (Coffman et al., 2011; Kelly & Strick, 2003; Middleton & Strick, 2001), reducing the influence the cerebellum has on cortical processing and in turn, the cortex is no longer able to rely on the cerebellum for support, and must recruit resources elsewhere.

### Limitations

While our findings provide a new insight into the role the cerebellum plays in cognitive processing, there are a few limitations worth noting. First, is electrode size. While a large portion of the literature has used the traditional 1×1 montage to modulate cerebellar function (Buch et al., 2017; Ferrucci et al., 2015), it is possible that stimulation occurred outside of the right cerebellum due to spread of the signal. Thus, we likely impacted a greater region of the cerebellum beyond the lateral posterior region we targeted, potentially including ventral cortical areas, in addition to the spinal cord and surrounding musculature. A second limitation is task difficulty, particularly during the Sternberg task. Though the current task seemed to be reasonably difficult in terms of memory load for young adults ^76^, accuracy levels across all groups were ∼ 90%. Therefore, there was not much room for modulation of task performance.

## Conclusion

Here using working memory and explicit motor sequence learning we demonstrated that cerebellar cathodal stimulation resulted in improved performance, and anodal stimulation hindered task performance. This effect of anodal stimulation also resulted in increased cortical activation, which we suggest is a compensatory mechanism due to the purported downregulation of the cerebellum after anodal stimulation. Specifically, when cerebellar output is degraded by anodal stimulation, the scaffolding effect the cerebellum provides is reduced, requiring more cortical activation to compensate for the reduced cerebellar output. This work has a potential to update existing models of aging and disease to include the cerebellum as a structure used to support cognitive processes, which has implications for remediation across clinical diagnoses.

## Acknowledgements

We would like to thank research assistants Sydney Eakin, Ivan Herrejon, and Sydney Cox for their help with data collection. The authors also acknowledge the Texas A&M University Brazos HPC cluster that contributed to the research reported here.

## Declarations

This research did not receive any specific grant from funding agencies in the public, commercial, or not-for-profit sectors. The authors have no relevant financial or non-financial interests to disclose. The authors do not have any conflicts of interest.

