## Supplemental Material for "Inhibitory Cerebellar Stimulation Increases Cortical Activation: Evidence for Cerebellar Scaffolding of Cortical Processing"

### Supplemental Results

#### Analysis of Order Effects

<u>Sequence Learning</u>

*RT.* In addition to the effects we already describe, we found a marginal effect of order (*p*=. 052), such that RT was faster when the sequence learning task was second (Supplemental Figure 1a). We also found a three-way interaction between stimulation, phase and order for RT (*p*< .001). Specifically, RT was better following anodal stimulation during early learning trials, when sequence was completed second (*p*=.040). No other effects were significant (*p*s> .09). We did not find stimulation by order (*p*=.842) or phase by order (*p*=.107) interactions. Broadly, this suggests that perhaps effects of tDCS on RT were different for sequence learning if the task was performed second. However, they are very specific and limited in their impact.

*Accuracy.* In addition to the effects we already describe, we found phase by order interaction (*p*< .001), such that accuracy was better for random trials when the sequence learning task was completed first (Supplemental Figure 1b). We also found a three-way interaction between stimulation, phase and order for accuracy (*p* = .042). Specifically, accuracy was better following sham stimulation during early learning trials, when sequence learning was completed second (*p*=.033). No other effects were significant (*p*s > .09). We did not find an effect of order (*p*=.393), or a stimulation by order (*p*=.747) interaction. This suggests that task order relative to stimulation had no impact on accuracy during the sequence learning task.

Together, there appear to be no impacts on stimulation order with respect to accuracy on the sequence learning task, and only small, limited effects on RT. The very limited nature of the RT effects suggest that these results are robust, and the impact of tDCS likely continued across the entire scanning period. Indeed, stimulation duration does modulate the time needed for cortical activation to return to baseline (Nitsche & Paulus, 2001). Specifically, 9 minutes of stimulation resulted in 30 minutes of effect and 13 minutes resulted in over 90 minutes of effect. Therefore, 20 minutes of stimulation should ensure the effect of stimulation was present for the duration of the scanning session, which was completed well within 90 minutes of stimulation. It should be worth noting that we might be experiencing an effect of comfort. That is, as participants get more familiar with responding to stimuli in a scanner, their performance improves.

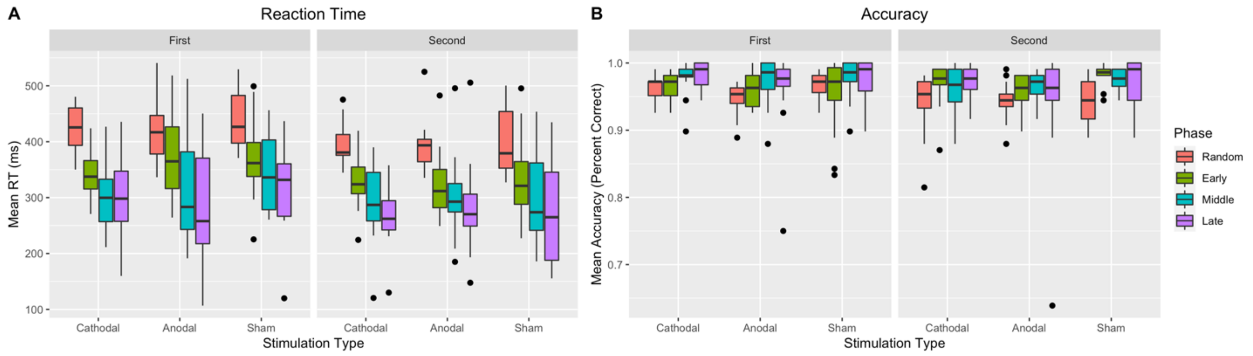


**Supplemental Figure 1**. Mean RT and accuracy for the sequence learning task by Phase, Region, Stimulation Condition, and Task Order. Dots indicate outliers. Whiskers represent the interquartile range.

<u>*Sternberg Task*</u>

*RT*. There were no effects of order onreaction time (*p*s>.090; Supplemental Figure 2a).

*Accuracy*. In addition to the effects we already describe, we found a three-way interaction between stimulation, load, and order for accuracy ($p < .001$; Supplemental Figure 2b). Specifically, accuracy was better following cathodal stimulation during high load trials, when Sternberg was completed second ($p$=.023). Also, accuracy was better following sham stimulation during high load, if Sternberg was completed first ($p$=.0167). No other effects were significant (*p*s > .214). We did not find an effect of order ($p$=.781), or stimulation by order ($p$=.241), or load by order ($p$=.474) interactions.

In general, there was some benefit to performing the task second, but it was limited to the cathodal condition or to sham, and very specific in terms of load. This could suggest a waning effect of stimulation, but certainly this impact was not broad. Further, given that one of the effects related to sham stimulation, it may again be the case that participants had an increased familiarity with making responses in the scanner environment.

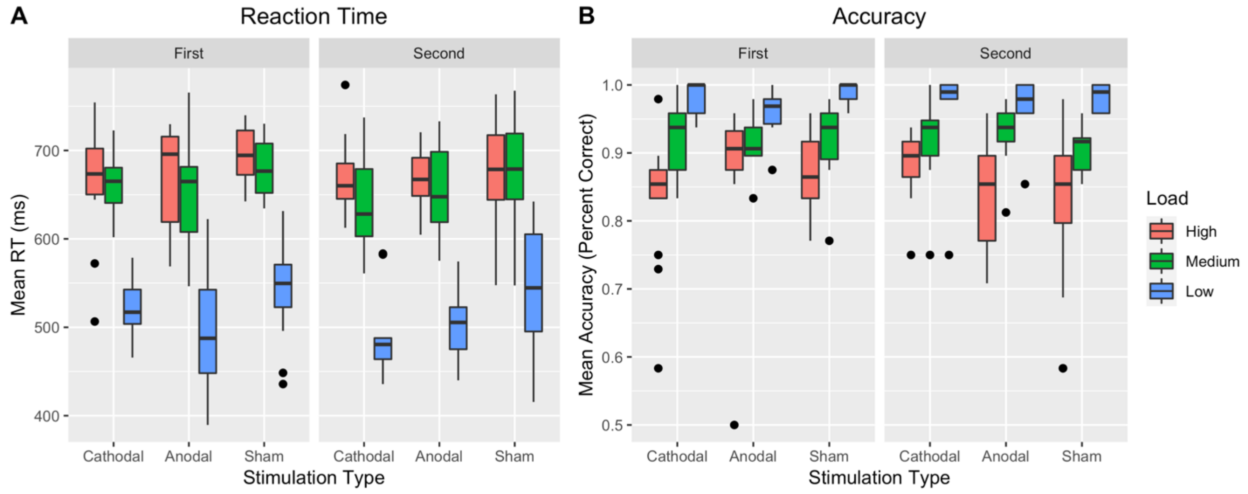


**Supplemental Figure 2**. Mean RT and accuracy for the Sternberg task by Load, Stimulation Condition, and Task Order. Dots indicate outliers. Whiskers represent the interquartile range.

**Sequence Learning**

<u>Behavioral Data</u>

**Supplemental Table 1**. Mean RT and accuracy for the sequence learning task by Phase, Region, and Stimulation Condition.

| | | Accuracy | | Reaction Time (ms) | |
|---|---|---|---|---|---|
| **Phase** | **Stimulation** | **Mean** | **SD** | **Mean** | **SD** |
| Early | Anodal | 0.96 | 0.20 | 350.74 | 132.50 |
| Early | Cathodal | 0.97 | 0.18 | 336.04 | 117.12 |
| Early | Sham | 0.96 | 0.18 | 359.10 | 126.63 |
| Late | Anodal | 0.95 | 0.23 | 293.23 | 135.62 |
| Late | Cathodal | 0.98 | 0.15 | 285.94 | 120.03 |
| Late | Sham | 0.97 | 0.17 | 311.98 | 131.05 |
| Middle | Anodal | 0.97 | 0.17 | 315.96 | 131.02 |
| Middle | Cathodal | 0.97 | 0.18 | 298.33 | 114.57 |
| Middle | Sham | 0.98 | 0.15 | 330.50 | 133.81 |
| Random | Anodal | 0.94 | 0.23 | 407.36 | 104.51 |
| Random | Cathodal | 0.95 | 0.22 | 409.86 | 103.70 |
| Random | Sham | 0.96 | 0.21 | 422.30 | 111.79 |

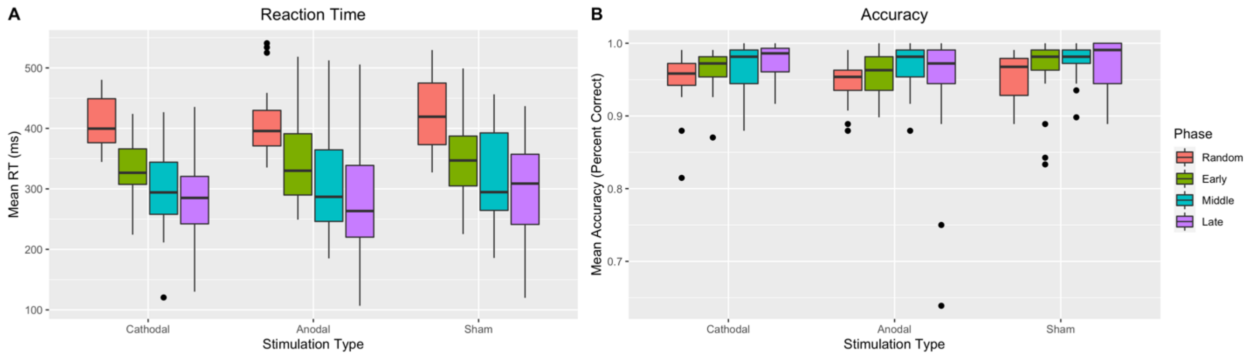


**Supplemental Figure 3**. Mean RT and accuracy for the sequence learning task by Phase, Region, and Stimulation Condition. Dots indicate outliers. Whiskers represent the interquartile range.

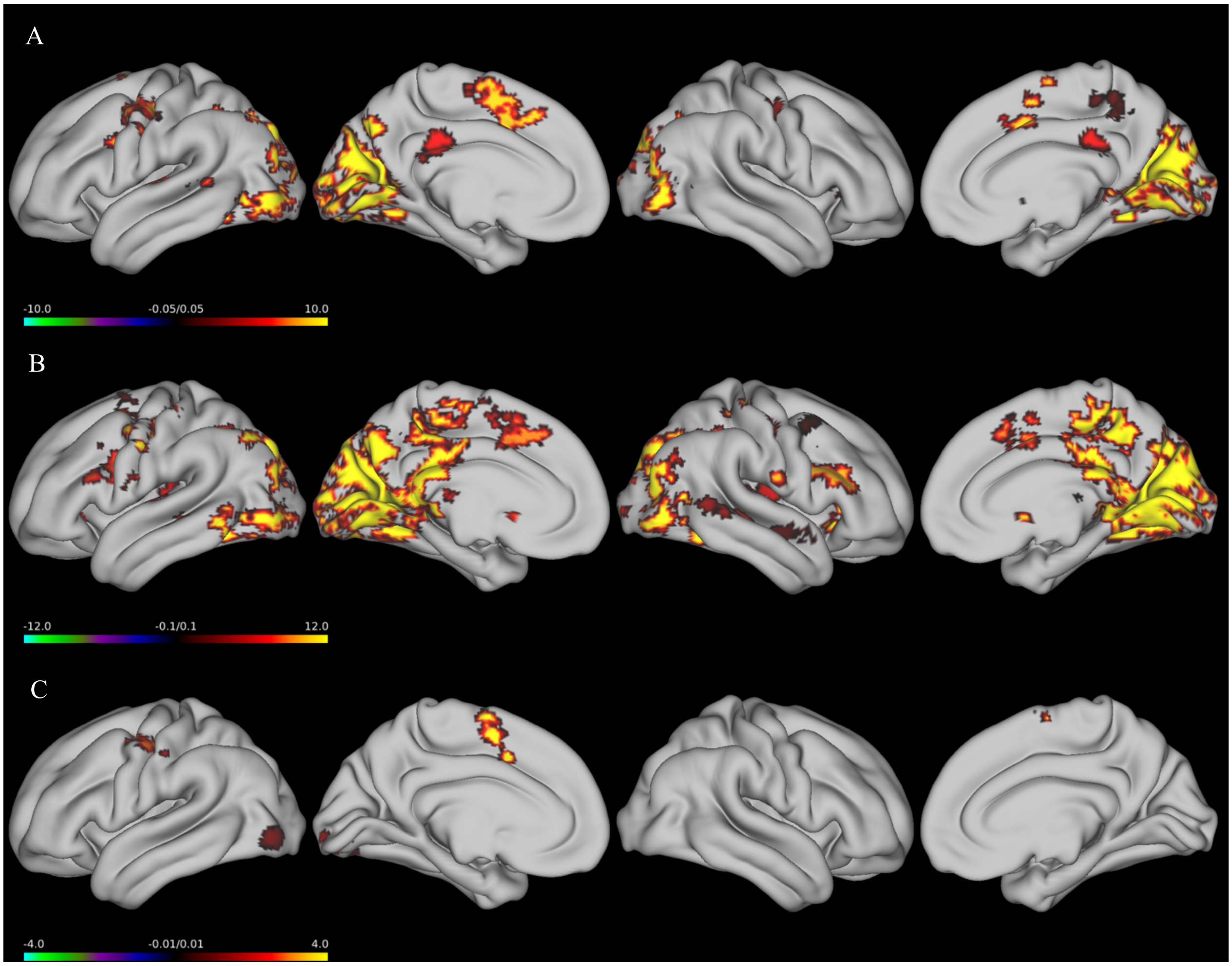


**Supplemental Figure 4**. Significant activations in the sham stimulation group during a Sequence learning task. (A) Sham activation during random trials. (B) Sham activation during sequence trials. (C) Sham activations for the sequence > random contrast.

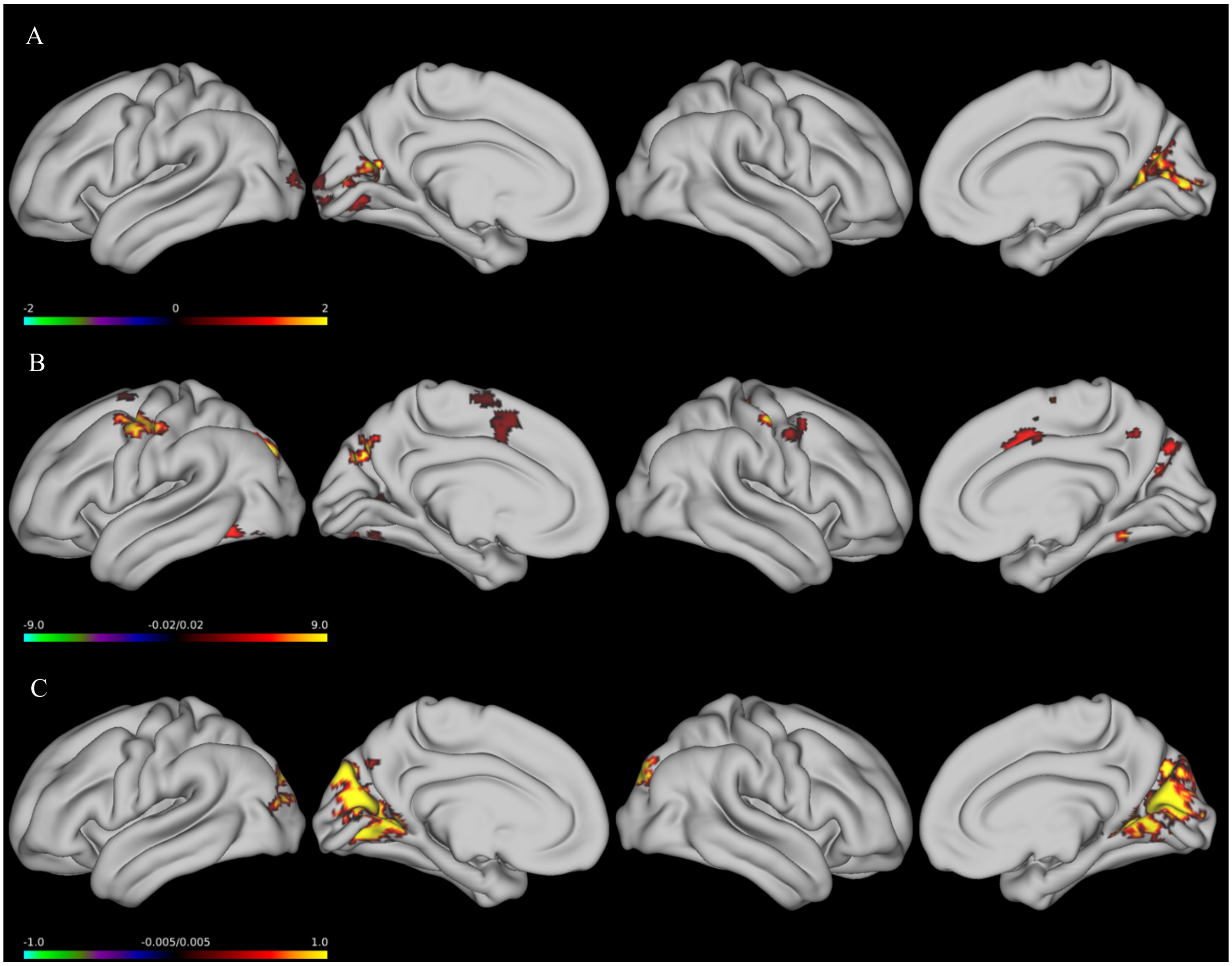


**Supplemental Figure 5**. Sham activations during early, middle and late learning phases. (A) Early learning; (B) Middle learning; (C) Late learning.

**Supplemental Table 2**. Significant clusters associated with explicit motor sequence learning following sham stimulation.

| Phase | Region | BA | Voxels | MNI Coordinates | | | Z |
|---|---|---|---|---|---|---|---|
| | | | | x | y | z | |
| Random | Left Calcarine fissure and surrounding cortex | Left-PrimVisual (17) | 18101 | -10 | -66 | 16 | 8.41 |
| | Left Lenticular nucleus, putamen | Left-Caudate (48) | 1388 | -16 | 18 | -2 | 6.72 |
| | Right Inferior frontal gyrus, triangular part | Right-BA9 | 732 | 50 | 26 | 24 | 5.93 |
| | Left Median cingulate and paracingulate gyri | Left-BA8 | 606 | -8 | 16 | 42 | 6.02 |
| | Left Inferior frontal gyrus, opercular part | Left-BA6 | 468 | -40 | 4 | 28 | 6.42 |
| | Left Superior temporal gyrus | Left-PrimAuditory (41) | 338 | -46 | -22 | 8 | 5.22 |
| | Right Heschl gyrus | Right-PrimSensory (1) | 298 | 48 | -12 | 10 | 5.93 |

| | | | | | | | |
|---|---|---|---|---|---|---|---|
| | Right Middle temporal gyrus | Right-BA21 | 254 | 68 | -40 | 4 | 4.98 |
| | Left Supplementary motor area | Left-BA6 | 194 | -4 | -2 | 64 | 4.92 |
| | Right Middle temporal gyrus | Right-BA22 | 150 | 48 | -14 | -12 | 4.96 |
| | Right Thalamus | Right-Thalamus (50) | 147 | 10 | -16 | 12 | 4.52 |
| | Right Middle frontal gyrus | Right-BA8 | 141 | 38 | 8 | 50 | 5.01 |
| Sequence | Right Cuneus | Right-BA31 | 8722 | 14 | -68 | 22 | 7.22 |
| | Left Supplementary motor area | Left-BA6 | 718 | -6 | 2 | 56 | 6.18 |
| | Left Postcentral gyrus | Left-PrimMotor (4) | 582 | -42 | -16 | 48 | 6.01 |
| | Left Heschl gyrus | Left-PrimMotor (4) | 291 | -54 | -10 | 10 | 4.9 |
| | Left Posterior cingulate gyrus | Left-BA23 | 276 | -4 | -30 | 30 | 4.78 |
| | Right Precental gyrus | Right-PrimMotor (4) | 258 | 36 | -18 | 44 | 4.93 |
| | Right Lenticular nucleus, putamen | Right-Putamen (49) | 249 | 26 | 12 | -6 | 4.97 |
| | Left Caudate nucleus | Left-Caudate (48) | 191 | -14 | 22 | -2 | 5.19 |
| | Right Paracentral lobule | Right-SensoryAssoc (5) | 158 | 12 | -40 | 56 | 4.69 |
| | Right Insula | Right-Insula (13) | 147 | 34 | 20 | 6 | 5.01 |
| Sequence> Random | Left Supplementary motor area | Left-BA6 | 196 | -6 | 2 | 56 | 6.25 |
| | Left Precental gyrus | Left-BA6 | 186 | -38 | -6 | 50 | 5.29 |
| | Left Lingual gyrus | Left-VisualAssoc (18) | 176 | -16 | -84 | -6 | 6.73 |
| | Left Inferior occipital gyrus | Left-VisualAssoc (18) | 123 | -38 | -86 | -6 | 6.01 |
| Early | Right Calcarine fissure and surrounding cortex | Right-PrimVisual (17) | 781 | 8 | -72 | 20 | 6.13 |
| | Left Lingual gyrus | Left-VisualAssoc (18) | 251 | -14 | -76 | -6 | 5.23 |
| Middle | Left Middle occipital gyrus | Left-BA39 | 553 | -28 | -72 | 36 | 5.75 |
| | Left Precental gyrus | Left-BA6 | 512 | -42 | -2 | 54 | 5.29 |
| | Right Precental gyrus | Right-PrimMotor (4) | 275 | 48 | -12 | 54 | 4.9 |
| | Right Fusiform gyrus | Right-BA19 | 235 | 30 | -58 | -4 | 5.24 |
| | Left Fusiform gyrus | Left-BA19 | 228 | -34 | -70 | -12 | 5.22 |
| | Right Superior occipital gyrus | Right-BA19 | 225 | 18 | -82 | 36 | 4.6 |
| | Right Median cingulate and paracingulate gyri | Right-BA32 | 146 | 8 | 4 | 40 | 4.99 |
| | Right Middle frontal gyrus | Right-BA6 | 139 | 34 | 6 | 52 | 4.99 |
| | Left Supplementary motor area | Right-BA6 | 129 | 0 | 10 | 54 | 4.87 |
| | Left Superior frontal gyrus, dorsolateral | Left-BA6 | 121 | -20 | 0 | 66 | 4.72 |
| | Left Calcarine fissure and surrounding cortex | Left-BA23 | 101 | -20 | -66 | 8 | 4.65 |
| Late | Left Cuneus | Left-VisualAssoc (18) | 3570 | -2 | -80 | 32 | 6.31 |

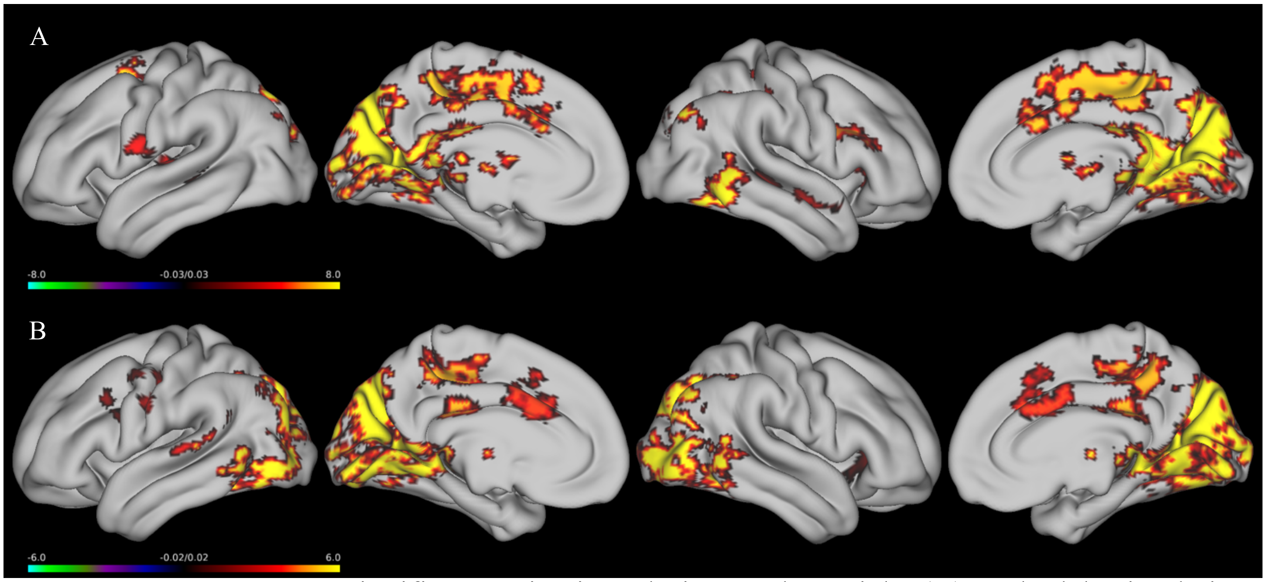


**Supplemental Figure 6**. Significant activations during random trials; (A) Cathodal Stimulation; (B) Anodal Stimulation.

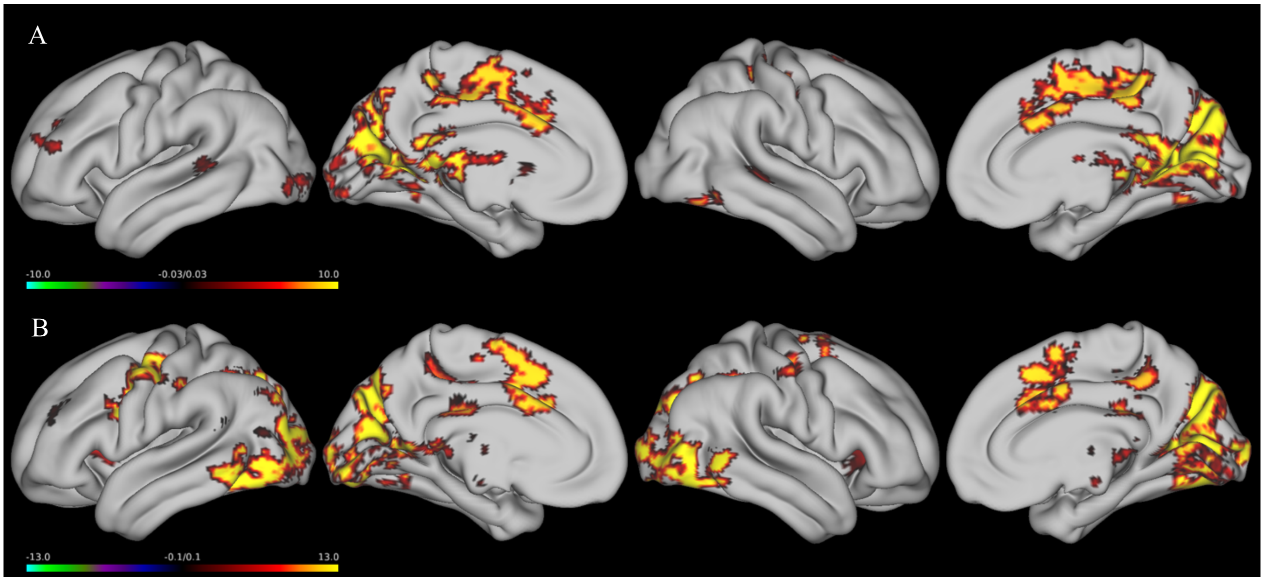


**Supplemental Figure 7**. Significant activations following sequence learning; (A) Cathodal Stimulation; (B) Anodal Stimulation.

**Supplemental Table 3**. Significant clusters associated with explicit motor sequence learning following active stimulation.

| Phase | Stimulation | Region | BA | Voxels | MNI Coordinates | | | Z |
|---|---|---|---|---|---|---|---|---|
| | | | | | x | y | z | |
| Random | Cathodal | Right Calcarine fissure and surrounding cortex | Right-VisualAssoc (18) | 11099 | 12 | -62 | 16 | 7.62 |
| | | Right Precuneus | Right-BA31 | 3550 | 4 | -42 | 54 | 6.13 |
| | | Right Inferior frontal gyrus, opercular part | Right-BA44 | 358 | 44 | 6 | 24 | 5.02 |
| | | Left Postcentral gyrus | Left-PrimMotor (4) | 271 | -58 | -4 | 20 | 4.95 |
| | | Right Temporal pole: superior temporal gyrus | Right-BA38 | 183 | 50 | 10 | -14 | 5.15 |
| | | Right Insula | Right-Insula (13) | 167 | 30 | 18 | 6 | 5.02 |
| | | Right Superior temporal gyrus | Right-BA22 | 160 | 46 | -26 | -2 | 5.31 |
| | | Left Middle temporal gyrus | Left-BA21 | 128 | -48 | -36 | 2 | 4.97 |
| | Anodal | Right Cuneus | Right-VisualAssoc (18) | 15095 | 6 | -76 | 32 | 9.97 |
| | | Right Precuneus | Right-BA31 | 1849 | 6 | -42 | 48 | 6.33 |
| | | Left Anterior cingulate and paracingulate gyri | Left-BA24 | 687 | -6 | 18 | 28 | 6.03 |
| | | Left Postcentral gyrus | Left-BA6 | 410 | -56 | -8 | 30 | 5.28 |
| | | Left Precental gyrus | Left-BA8 | 135 | -48 | 12 | 36 | 5.22 |
| | | Right Insula | Right-Insula (13) | 115 | 38 | 20 | -8 | 5.7 |
| Sequence | Cathodal | Left Calcarine fissure and surrounding cortex | Left-PrimVisual (17) | 5009 | -4 | -70 | 12 | 6.6 |
| | | Right Supplementary motor area | Right-BA6 | 3008 | 2 | -2 | 58 | 5.32 |
| | | Right Inferior temporal gyrus | Right-Fusiform (37) | 254 | 50 | -58 | -12 | 5.22 |
| | | Left Middle occipital gyrus | Left-VisualAssoc (18) | 244 | -20 | -100 | -2 | 5.12 |
| | | Left Lenticular nucleus, pallidum | Left-Caudate (48) | 200 | -8 | 4 | 2 | 4.92 |
| | | Left Middle frontal gyrus | Left-BA10 | 189 | -36 | 44 | 18 | 5.67 |
| | | Left Middle temporal gyrus | Left-BA22 | 140 | -48 | -38 | 6 | 5.14 |
| | | Right Middle temporal gyrus | Right-BA21 | 122 | 52 | -36 | 2 | 4.66 |
| | | Right Olfactory cortex | Right-Putamen (49) | 107 | 18 | 12 | -12 | 4.91 |
| | | Left Middle occipital gyrus | Left-BA39 | 106 | -28 | -74 | 32 | 4.78 |
| | Anodal | Right Cuneus | Right-BA19 | 12038 | 10 | -78 | 36 | 7.7 |
| | | Left Supplementary motor area | Left-BA6 | 1438 | -4 | 14 | 46 | 6.07 |
| | | Left Median cingulate and paracingulate gyri | Left-BA23 | 520 | -6 | -24 | 30 | 5.32 |
| | | Right Precental gyrus | Right-PrimMotor (4) | 277 | 44 | -16 | 52 | 5.33 |

| | | | | | | | |
|---|---|---|---|---|---|---|---|
| | Left Median cingulate and paracingulate gyri | Left-BA31 | 223 | -14 | -36 | 46 | 4.93 |
| | Right Thalamus | Right-Thalamus (50) | 218 | 8 | -18 | 4 | 5.7 |
| | Left Insula | Left-Insula (13) | 170 | -38 | 14 | 4 | 6.04 |
| | Left Lenticular nucleus, putamen | Left-Caudate (48) | 166 | -14 | 14 | -4 | 4.85 |
| | Left Thalamus | Left-Thalamus (50) | 164 | -8 | -18 | 8 | 5.43 |
| | Right Insula | Right-BA45 | 143 | 34 | 26 | 6 | 5.23 |
| | Right Lenticular nucleus, putamen | Right-Putamen (49) | 130 | 18 | 14 | -6 | 4.94 |
| | Left Middle temporal gyrus | Left-BA39 | 124 | -62 | -52 | 18 | 5.82 |
| | Left Middle frontal gyrus | Left-BA10 | 111 | -30 | 46 | 24 | 5.76 |

**Sternberg Task**

Behavioral Data

**Supplemental Table 4**. Mean RT and accuracy for the Sternberg task by Load, Region, and Stimulation Condition.

| | | **Reaction Time (ms)** | | **Accuracy** | |
|---|---|---|---|---|---|
| Load | Stimulation | Mean | SD | Mean | SD |
| Low | Anodal | 500.97 | 131.49 | 0.97 | 0.18 |
| Low | Cathodal | 506.79 | 121.54 | 0.98 | 0.15 |
| Low | Sham | 541.77 | 127.06 | 0.99 | 0.11 |
| Medium | Anodal | 652.43 | 197.95 | 0.93 | 0.26 |
| Medium | Cathodal | 649.92 | 198.77 | 0.92 | 0.27 |
| Medium | Sham | 676.84 | 195.10 | 0.91 | 0.28 |
| High | Anodal | 667.76 | 247.57 | 0.85 | 0.36 |
| High | Cathodal | 666.20 | 246.53 | 0.85 | 0.36 |
| High | Sham | 686.15 | 262.03 | 0.84 | 0.36 |

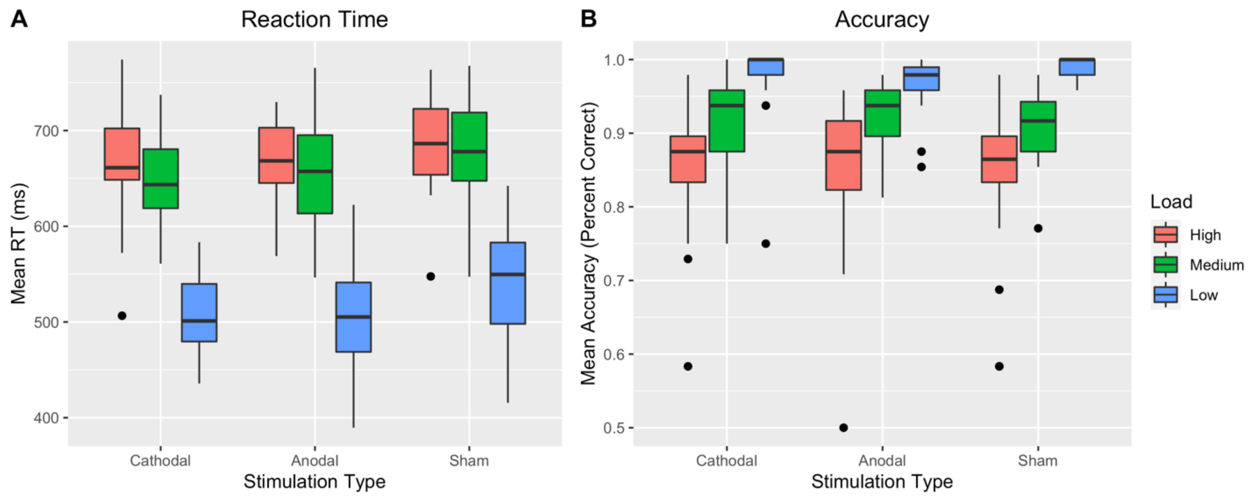


**Supplemental Figure 8**. Mean RT and accuracy for the Sternberg task by Load and Stimulation Condition. Dots indicate outliers. Whiskers represent the interquartile range. Both anodal and cathodal stimulation improved reaction time.

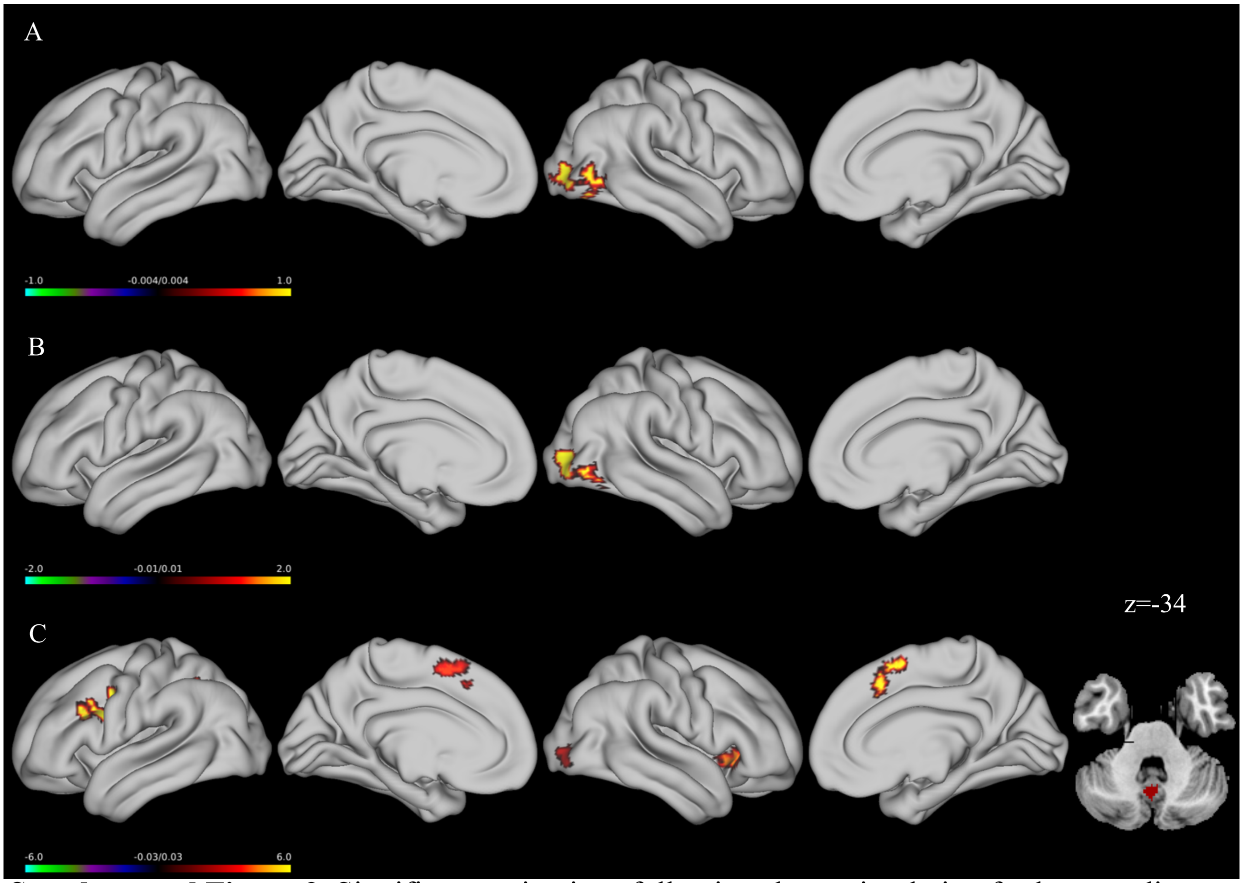

**Supplemental Figure 9**. Significant activations following sham stimulation for low, medium and high load; (A) Low Load; (B)Medium Load; (C) High Load.

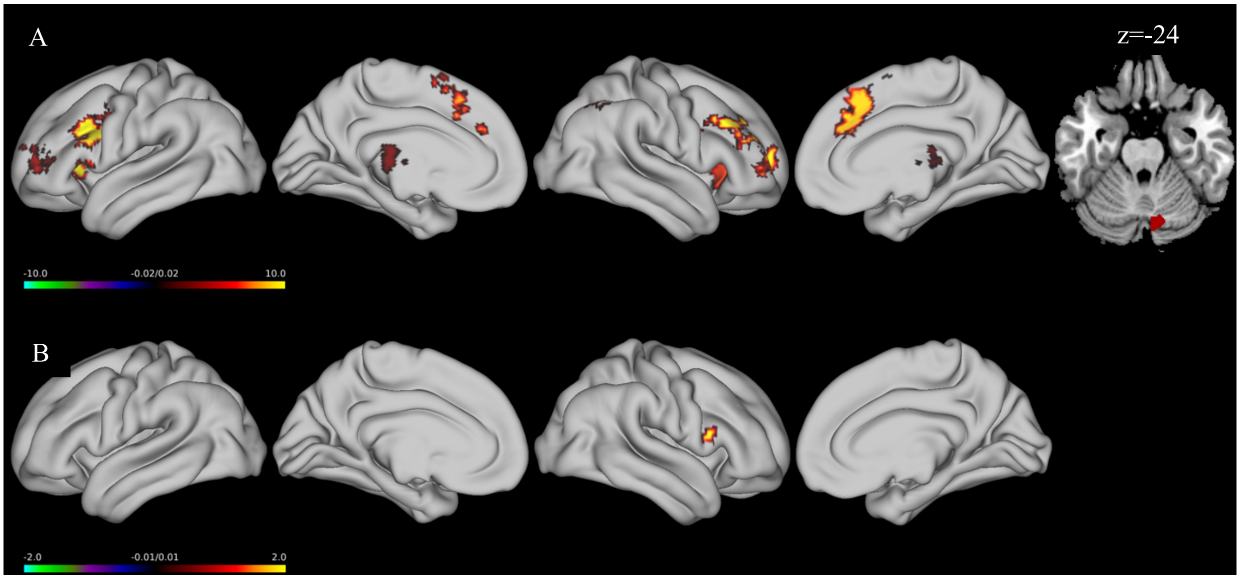

**Supplemental Figure 10**. Significant activations following sham stimulation for low, medium and high load contrasts; (A) Greater activation following high load compared to low load; (B) Greater activation during high load compared to medium load.

**Supplemental Table 5**. Significant clusters following sham stimulation during a Sternberg task.

| Load | Region | BA | Voxels | MNI Coordinates | | | Z |
|---|---|---|---|---|---|---|---|
| | | | | x | y | z | |
| Low | Right Middle occipital gyrus | Right-VisualAssoc (18) | 445 | 34 | -88 | 4 | 5.27 |
| Medium | Right Inferior occipital gyrus | Right-VisualAssoc (18) | 497 | 38 | -86 | -4 | 6.28 |
| | Left Inferior parietal, but supramarginal and angular gyri | Left-BA40 | 109 | -40 | -46 | 44 | 5.67 |
| High | Left Inferior frontal gyrus, opercular part | Left-BA44 | 448 | -44 | 10 | 22 | 5.26 |
| | Left Inferior parietal, but supramarginal and angular gyri | Left-BA40 | 305 | -40 | -44 | 42 | 5.83 |
| | Left Supplementary motor area | Left-BA6 | 247 | -4 | 6 | 56 | 5.67 |
| | Right Insula | Right-Insula (13) | 170 | 42 | 16 | -8 | 5.6 |
| | Right Inferior occipital gyrus | Right-VisualAssoc (18) | 123 | 36 | -86 | 0 | 5.29 |
| | Vermis VIII, cerebellum | Left-VisualAssoc (18) | 107 | -2 | -64 | -36 | 5.35 |
| High > Low | Left Inferior frontal gyrus, triangular part | Left-BA9 | 849 | -46 | 22 | 30 | 7.18 |
| | Right Inferior frontal gyrus, triangular part | Right-BA9 | 678 | 46 | 30 | 32 | 8.26 |
| | Left Supplementary motor area | Left-BA8 | 504 | -2 | 20 | 52 | 6.07 |
| | Left Inferior parietal, but supramarginal and angular gyri | Left-BA39 | 332 | -34 | -60 | 46 | 6.58 |
| | Right Insula | Right-Insula (13) | 267 | 32 | 24 | -2 | 6.14 |
| | Left Middle frontal gyrus | Left-BA10 | 194 | -30 | 54 | 12 | 6.5 |
| | Right Inferior parietal, but supramarginal and angular gyri | Right-BA40 | 159 | 44 | -44 | 42 | 6.2 |
| | Left Thalamus | Left-Thalamus (50) | 144 | -4 | -24 | 8 | 5.08 |
| | Right Thalamus | Right-Thalamus (50) | 121 | 8 | -10 | 10 | 5.33 |
| | Right Crus I, cerebellum | Right-VisualAssoc (18) | 106 | 10 | -72 | -26 | 6.07 |
| High > Medium | Right Inferior frontal gyrus, opercular part | Right-BA44 | 211 | 38 | 14 | 10 | 5.58 |
| | Left Inferior parietal, but supramarginal and angular gyri | Left-BA40 | 118 | -36 | -42 | 34 | 5.58 |

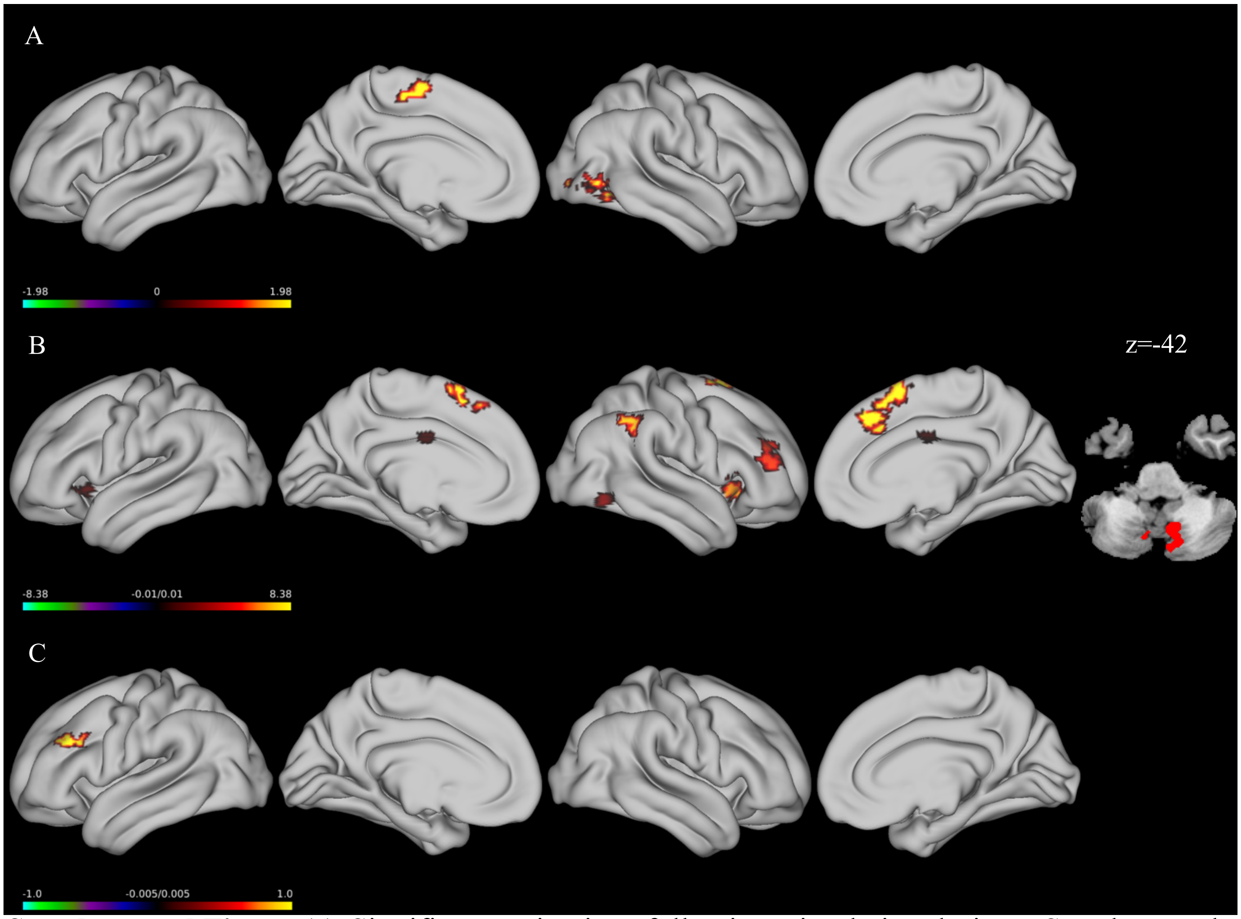


**Supplemental Figure 11**. Significant activations following stimulation during a Sternberg task. (A) Significant activations following cathodal stimulation during low load trials; (B) Significant activations following cathodal stimulation during high load trials; (C) Significant activations following anodal stimulation during high load trials.

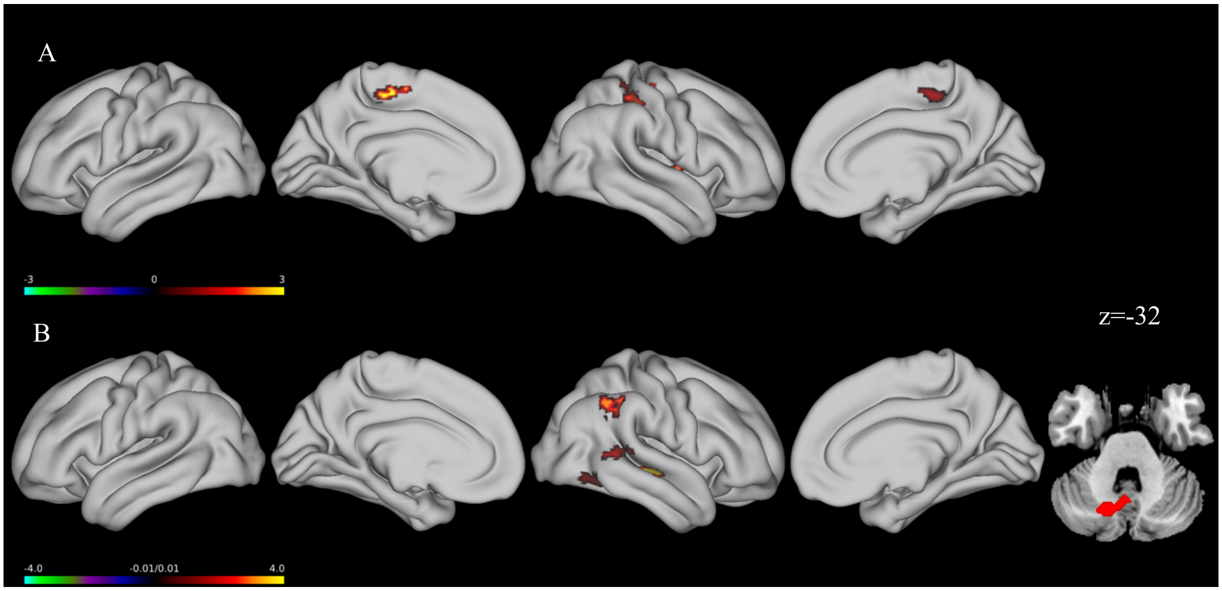

**Supplemental Figure 12**. Stimulation group differences. (A) Cathodal>Anodal during low load trials. (B) Cathodal>Anodal during the high load condition.

**Supplemental Table 6**. Significant clusters following stimulation during a Sternberg task.

| Load | Stimulation | Region | BA | Voxels | MNI Coordinates x | y | z | Z |
|---|---|---|---|---|---|---|---|---|
| Low | Cathodal | Right Middle temporal gyrus | Right-BA19 | 215 | 42 | -72 | 0 | 5.16 |
| | | Left Supplementary motor area | Left-BA6 | 108 | -4 | -16 | 58 | 4.76 |
| | Cathodal > Anodal | Right Lenticular nucleus, putamen | Right-Putamen (49) | 287 | 30 | -14 | 4 | 4.74 |
| | | Right Inferior parietal, but supramarginal and angular gyri | Right-PrimSensory (1) | 266 | 40 | -36 | 54 | 4.82 |
| | | Left Paracentral lobule | Right-BA6 | 174 | 0 | -24 | 54 | 4.92 |
| High | Cathodal | Right Superior frontal gyrus, medial | Right-BA8 | 472 | 6 | 34 | 42 | 5.32 |
| | | Right Angular gyrus | Right-BA39 | 280 | 36 | -54 | 38 | 5.15 |
| | | Right Insula | Right-Insula (13) | 258 | 36 | 22 | -6 | 6.07 |
| | | Right Middle frontal gyrus | Right-BA46 | 232 | 38 | 42 | 8 | 5.05 |
| | | Right Lobule VIIb, cerebellum | Right-VisualAssoc (18) | 221 | 6 | -76 | -44 | 5.04 |
| | | Right Inferior parietal, but supramarginal and angular gyri | Right-BA7 | 164 | 48 | -46 | 56 | 5.45 |
| | | Right Inferior temporal gyrus | Right-Fusiform (37) | 147 | 48 | -62 | -12 | 5.72 |
| | | Left Temporal pole: superior temporal gyrus | Left-BA47 | 120 | -48 | 16 | -10 | 5.03 |
| | | Right Median cingulate and paracingulate gyri | Left-BA24 | 115 | -2 | -6 | 30 | 4.97 |
| | Anodal | Left Inferior frontal gyrus, triangular part | Left-BA9 | 128 | -46 | 30 | 28 | 4.73 |
| | Cathodal > Anodal | Left Lobule VIII, cerebellum | Left-VisualAssoc (18) | 260 | -12 | -66 | -36 | 4.64 |
| | | Right Superior temporal gyrus | Right-PrimAuditory (41) | 159 | 52 | -16 | 2 | 4.73 |
| | | Right Inferior parietal, but supramarginal and angular gyri | Right-BA39 | 147 | 52 | -52 | 40 | 4.54 |
| | | Right Middle temporal gyrus | Right-BA22 | 139 | 54 | -42 | 8 | 4.73 |
| | | Right Inferior temporal gyrus | Right-Fusiform (37) | 116 | 50 | -56 | -10 | 5.53 |
